# The fatty acid synthesis pathway is a checkpoint for lipoteichoic acid synthesis in *Staphylococcus aureus*

**DOI:** 10.64898/2026.04.06.715823

**Authors:** Paprapach Wongdontree, Clara Louche, Julien Dairou, Vincent Leguillier, Christine Péchoux, Bastien Prost, Myriam Gominet, Karine Gloux, Patrick Trieu-Cuot, Audrey Solgadi, Alexandra Gruss, Jamila Anba-Mondoloni

## Abstract

Bacterial membranes comprise diverse lipids whose proportions vary according to environmental conditions. How cells direct lipid flux toward specific products remains unclear. We address this question in the human pathogen *Staphylococcus aureus*, where multiple lipid products compete for a common precursor, the major phospholipid, phosphatidylglycerol (PG). One product, lipoteichoic acid (LTA), is essential for cell division, envelope homeostasis, and virulence. Lipids and metabolites were quantified to identify factors that prioritize LTA synthesis over the other PG-derived products. We identify upstream fatty acid synthesis (FASII) pathway as a key control point for LTA production. Inhibition of FASII by antibiotics or gene inactivation causes LTA depletion. FASII inhibition similarly affects LTA in *Streptococcus agalactiae*, suggesting conservation of this LTA control strategy. Changes in membrane fatty acids do not account for LTA depletion. Instead, we show that FASII inhibition causes a drop in intracellular glycerophosphate (GroP), a precursor for both PG and LTA. Under these conditions of GroP limitation, PG flux favors production of a non-GroP lipid, cardiolipin. Moreover, combined inhibition of FASII and WTA blocks *S. aureus* growth, confirming the lethality of depleting LTA and WTA simultaneously. This study resolves how *S. aureus* manages phospholipid flux, by prioritizing the synthesis of GroP-rich LTA or of non-GroP-containing lipids according to FASII-controlled GroP availability.

## Introduction

Phospholipids are primary components of most bacterial membranes, and are essential for cell integrity. In *Staphylococcus aureus* and other Gram-positive pathogens, phospholipids are building blocks for membrane-anchored structures. Among them, lipoteichoic acid (LTA) represents ∼12 mole % of total membrane outer leaflet lipids [1]. *S. aureus* LTA comprises a glycerophosphate (GroP) polymer (∼25-mer) usually anchored to the membrane *via* a di-glucosyl diacylglycerol (DG-DAG) lipid [2]. In *S. aureus*, about three quarters of GroP polymers are decorated with D-alanines, which reduce the overall negative charge of the polymer and contribute to antimicrobial peptide resistance [1–5] [6–8]. The dedicated enzymes responsible for LTA synthesis, control of the GroP polymer tail length, and turnover of the GroP donor lipid in *S. aureus* are well characterized [9–14].

LTA is implicated in basic bacterial processes of cell division, autolysis, and antimicrobial resistance, and also mediates bacteria-host interactions, which all relate to its physical properties [1, 15–17]. It is proposed to produce a stiff repulsive brush, which together with wall teichoic acid (WTA), creates a barrier contributing to turgor pressure maintenance on the cell exterior. Bacterial septation is suggested to be coordinated by LTA binding to autolysin, while the structure and charge of LTA outside the membrane would offer protection against the pressure gradient inside and outside the cell [16, 18, 19]. Both features of LTA could contribute to its roles in cell division.

While *ltaS* is essential for LTA synthesis, genes mediating polymer anchoring, i.e., *ypfP* (also called *ugtP*) encoding diacylglycerol glucosyltransferase, and *ltaA*, encoding the DG-DAG anchor flippase, are not [20]. Both *ypfP* and *ltaA* mutants produce longer GroP polymers, and the polymer is suggested to use alternative anchors (i.e., phosphatidylglycerol (PG) or lysyl-PG [20, 21]; in one exception, the *ypfP* mutant derived from SA113 was LTA-negative [22]). In addition, the CozEb protein is involved in flipping the glycolipid anchor, and mutants also produce longer polymers [23]. Overall, these mutants produce comparable amounts of LTA as the wild type (WT) [23, 24].

Despite its essential roles, LTA is reportedly dispensable or deleterious in specific conditions or mutant backgrounds. Growth of an *ltaS* mutant (defective for the LTA synthase LtaS) is restored in high osmolarity or low temperature [16, 18, 25, 26]. Growth defects in *clpX* [27], *pgl* [28], and *cshA* [29] are rescued by mutations in *ltaS*; LTA was absent or reduced in these mutants. Conversely, *ltaS* mutant growth was rescued by various suppressor mutations (*cozEb*, *sgtB*, *mazEF*, and *clpX*; [30]). Interestingly, expression of LtaS, which is essential for LTA synthesis, decreases sharply in stationary phase [31]. These findings predict that regulation of LTA expression may be multifactorial and condition-dependent, and that LTA is not always required for growth.

Remarkably, LTA synthesis is only one of 4 possible pathways using PG as a substrate, suggesting that these pathways are in competition, creating a ‘metabolic fork dilemma’ (**Fig. 1**). The factors that determine which end-product is favored are unknown. Solving this dilemma may be complex, as LTA synthesis builds upon upstream pathways that provide PG precursors, namely for fatty acid (FA) and GroP synthesis, to generate the LTA lipid anchor and GroP polymer. Moreover, the FA synthesis pathway (FASII) in *S. aureus* and numerous Bacillota is dispensible and growth is fully compensated by environmental FAs [32–35]. The shift from FASII to exogenous FA utilization (called here ‘FASII^bypass^’) is accompanied by marked changes in lipid metabolism and protein expression: these include energy savings by not using FASII, and reverse directionality of the glycerol-3-phosphate acyltransferase PlsX compared to FASII (**Supplementary Fig. S1A** and **S1B**) [35]. Here we established a causal link between FASII activity and LTA synthesis in *S. aureus*, and give evidence for the generality of this control in other species. Our findings suggest a simple mechanism for shunting PG towards LTA synthesis PG, based on availability of GroP, a metabolite shared by both products and controlled by FASII. LTA depletion during FASII bypass leads to an increase in WTA, opening perspectives for a bitherapy treatment as shown in a proof of concept demonstration.

**Figure 1.**
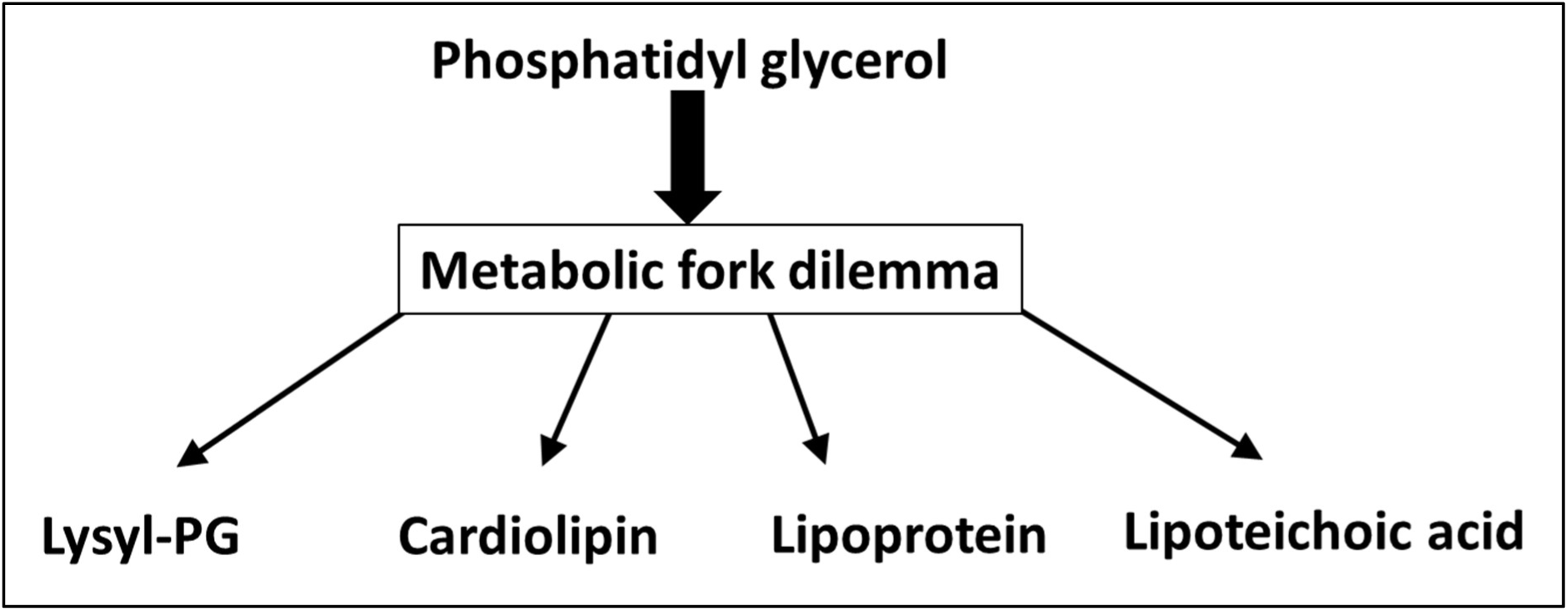
The metabolic dilemma of producing four possible lipids from one substrate. Phosphatidylglycerol (PG) gives rise to four alternative lipid products. The factors that may favor the flux towards one of these pathways are unknown.

## Results

### FASII inhibition induces *S. aureus* cell envelope changes during adaptation

*S. aureus* adapts to FASII inhibition (FASII^bypass^) by using exclusively environmental FAs to produce membrane phospholipids. FASII^bypass^ is a non-mutational event, but may also occur upon mutation of FASII initiation genes *acc* and/or *fabD* [33, 34, 36, 37]. Bacteria adapted to an anti-FASII proliferate exponentially after an initial latency phase (6 to 10 h according to conditions) (**Supplementary Fig. S1B**), and undergo profound protein expression changes [33, 35]. *S. aureus* WT USA300 JE2 strain (called JE2) was examined by transmission electron microscopy during adaptation to the anti-FASII (FabI inhibitor AFN-1252, 0.5 µg/ml [38]) (**Fig. 2A**). At 6 h post-anti-FASII treatment, prior to full adaptation, bacteria show a pronounced transient increase in envelope thickness (∼28.6 nm in non-treated bacteria, to 42.6 nm). Once adapted (at 10 h), envelope thickness is slightly greater than that of non-treated bacteria (∼29.7 nm, p=0.04), and cell division septa appear normally positioned. Overall, membrane contours were more irregular in the FASII^bypass^ bacteria (**Fig. 2A** and inset, **Supplementary Fig. S2**). Alterations in membrane integrity were observed in FASII^bypass^, as retention of a membrane-permeable drug, ethidium bromide (EtBr), was greater in FASII^bypass^ than in non-treated bacteria, particularly in stationary phase (**Fig. 2B**).

**Figure 2.**
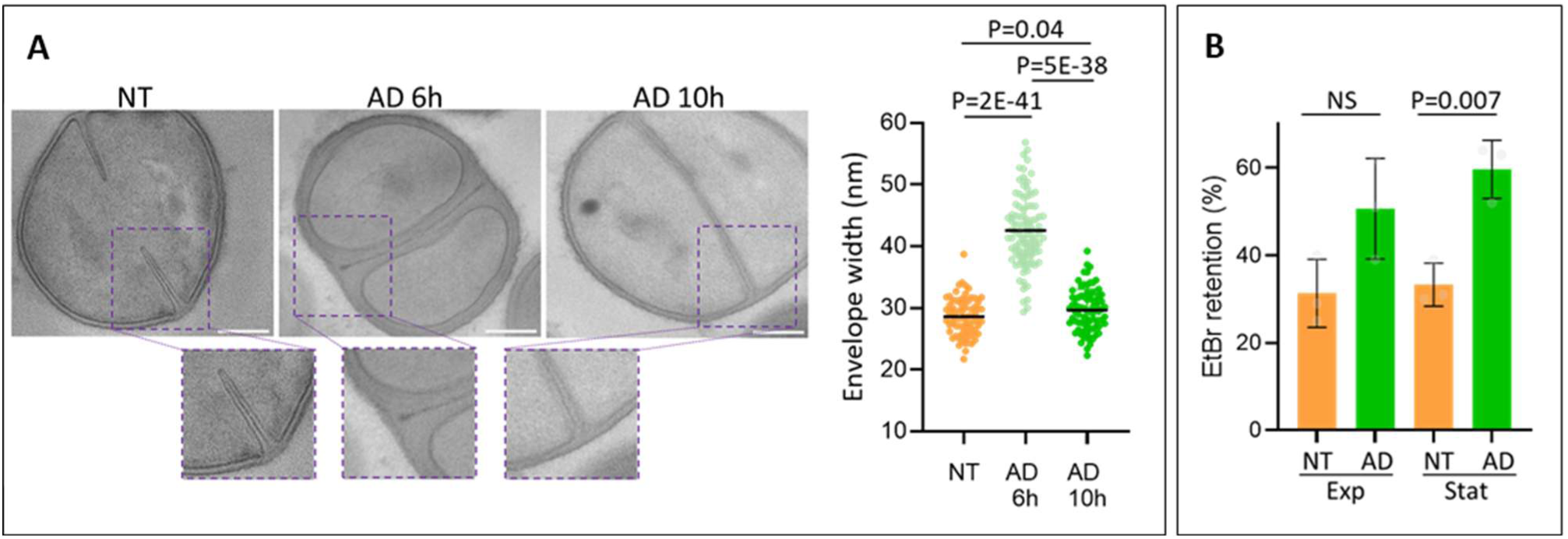
Morphological changes in the *S. aureus* envelope accompany anti-FASII adaptation. The JE2 strain was grown in SerFA (BHI containing 10% serum and an equimolar mixture of 3 FAs) or in SerFA supplemented with the anti-FASII (AFN-1252, 0.5 µg/ml; SerFA-AFN). **A.** Transmission electron microscopy: left, exponential (3 h) growth of non-treated *S. aureus* in SerFA to OD_600_ = ∼3; middle, 6 h growth post anti-FASII-treatment; right, 10 h post-anti-FASII-treatment, OD_600_ = ∼3. White bar, 200 nm. A zoom of the surface within the dotted box highlights morphological differences in septal regions. Envelope thickness in the three conditions are shown in the graph at right. P-values were determined using a two-sided T-test based on 73, 89, and 60 measurements respectively on at least 15 individual bacteria (Fig. 2A Source data). Additional images are in **Supplementary Fig. S2**. **B.** Ethidium bromide (EtBr) retention was compared in non-treated (NT) and anti-FASII-adapted (AD) *S. aureus* in exponential (exp, N=3) and stationary (stat; N=6) phase cultures. EtBr retention is presented at 20 minutes; see Fig. 2B Source data for the full data set. P-values were determined using a two-sided T-test on biological triplicates for exponential and six replicates for stationary cultures. NS, non-significant.

A recent proteomics analysis of *S. aureus* responses to the anti-FASII triclosan [33, 35] was rescreened to detect alterations in envelope biosynthesis functions (**Supplementary Fig. S3**). Notably, the diacylglycerol glucosyltransferase YpfP (SAUSA300_0918, also called UgtP), which synthesizes the LTA glycolipid diglucose anchor [11], was detected in non-treated bacteria, but dropped below detection levels upon anti-FASII treatment. *ypfP* is not essential for growth or LTA formation, but GroP polymers may be longer in the mutant than in WT strains [10]. The above observations were suggestive of envelope structural differences between the two bacterial states, and motivated us to evaluate LTA production in anti-FASII-adapted *S. aureus*.

### FASII arrest leads to LTA depletion

LTA abundance was compared in the *S. aureus* JE2 non-treated and FASII^bypass^ cultures by immunoblotting with LTA-specific antibody (Clone 55; [39]). LTA was readily detected in non-treated bacteria, and at 6 h post-AFN-1252 treatment. However, LTA levels diminished at 10 h post treatment (**Fig. 3A**). Upon dilution of FASII^bypass^ cultures, regrowth occurred without a lag phase and LTA remained depleted, indicating that anti-FASII adaptation exerted a long-term effect; FA profiles remained exogenous in this condition (**Supplementary Fig. S4**). LTA levels were also depleted in the WT strain treated by another FASII inhibitor, platensimycin, which targets FabF (3-oxoacyl-(acyl-carrier-protein) synthase II) [34, 40, 41], showing that the observed effect was not antibiotic-specific (**Fig. 3B**). An FA-auxotroph *fabD* deletion mutant, which relies on FASII^bypass^ for growth, similarly led to LTA depletion (**Fig. 3A**, right). Strain specificity of the LTA response to FASII arrest was ruled out, as LTA loss upon FASII^bypass^ was observed in a different *S. aureus* lineage, NCTC_8325, shown for strains RN4220-R and HG1-R (tested with AFN-1252; **Supplementary Fig. S5**). Thus all tested situations of *S. aureus* FASII^bypass^ and growth compensation by exogenous FAs cause LTA depletion.

**Figure 3.**
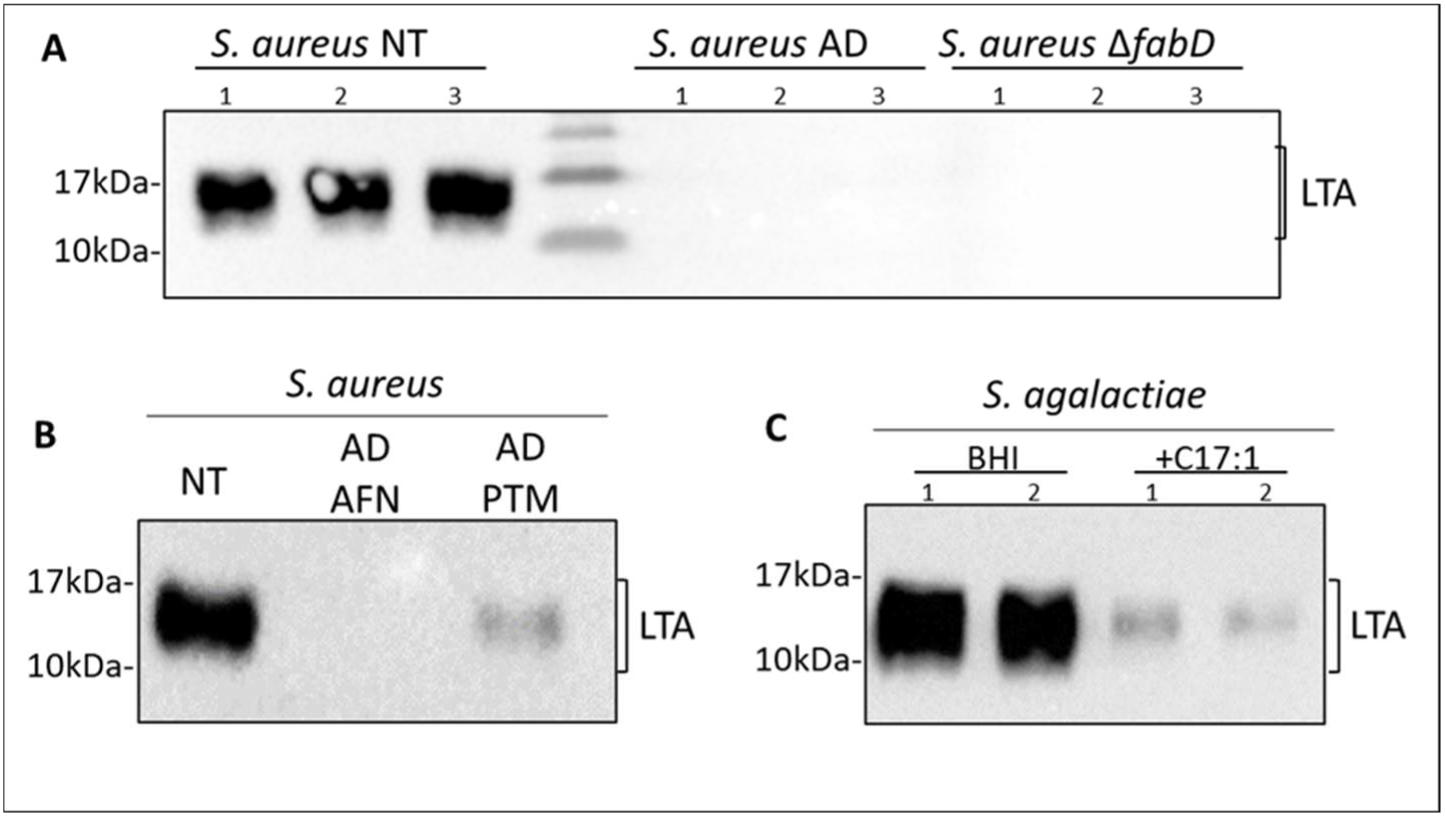
FASII^bypass^ leads to LTA depletion in *S. aureus* and *S. agalactiae*. *S. aureus* and numerous streptococci compensate FASII inhibition, deletion, or repression, by incorporating exogenous FAs in membranes [32, 33, 43, 65]**. A.** Antibiotic- or *fabD-*mutation- generated FASII inhibition leads to LTA depletion in *S. aureus*. WT JE2 was grown in SerFA without (NT, non-treated) or with FabI inhibitor AFN-1252 (AD, anti-FASII-adapted) [38]. Δ*fabD* is a FASII auxotroph ([34, 40]), and was grown in SerFA (N=3). **B.** Both anti-FabI AFN-1252 (AFN) and anti-FabF platensimycin (PTM, [41]) lead to LTA depletion in *S. aureus* (N=3). **C.** Inhibition of FASII by exogenous FAs in the streptococcus species *S. agalactiae* leads to LTA depletion. The *S. agalactiae* FASII pathway is repressed by FabT when bound to exogenous FAs [32]. Here, *S. agalactiae* strain NEM316 was grown in BHI plus 0.025% FA-free bovine serum albumin without or with C17:1 (100 µM); exogenous FA addition represses FASII but allows bacterial growth. LTA was monitored by immunoblotting using anti-LTA antibody (N=5). Samples were taken from exponential phase OD_600_ = ∼2-3 (**A** and **B**) and OD_600_ = ∼1 (**C**).

The potential generality of connecting FASII activity to LTA production was examined using the major pathogen *Streptococcus agalactiae*, which produces LTA with a GroP polymer similar to that of *S. aureus* [42]. In this and other streptococci, FASII enzymes are feedback-inhibited *via* exogenous FA-mediated activation of FabT, which represses FASII by forming an acyl-ACP-FabT complex. The consequence is that streptococcal membrane phospholipids exclusively comprise exogenous FAs whenever bacteria grow in lipid-containing environments [32, 43]. LTA was depleted when *S. agalactiae* NEM316 was grown in medium supplemented with a single FA, here C17:1*cis* (**Fig. 3C**). We conclude that blocking FASII causes LTA depletion and can be generalized to different *S. aureus* lineages, and to a streptococcus species.

### LTA levels are not restored by LtaS expression

Of the three main LTA synthesis enzymes, YpfP, the antiporter flippase LtaA (SAUSA300_0917, encoded adjacently to *ypfP*), and the LTA synthase LtaS, which catalyzes GroP polymerization (SAUSA300_0703), the first two are not essential and may be substituted by other *S. aureus* functions, although LTA gel migration and LTA localization are affected [39]. During FASII^bypass^, detected traces of LTA migrate like the non-treated samples (e.g., **Fig. 3**) and cell morphology appears normal (**Supplementary Fig. S2**), suggesting that decreased YpfP levels are not the cause of LTA depletion. We focused on LtaS as potentially explaining LTA depletion, as it catalyzes GroP polymerization, and is essential for LTA synthesis [44]. We first evaluated LtaS status in non-treated and FASII^bypass^ *S. aureus* by comparing sensitivity to Congo Red, an LtaS inhibitor [45]. Compared to non-treated *S. aureus*, Congo Red resistance of anti-FASII-adapted bacteria was ∼10-fold increased (**Fig. 4A**), suggesting that LtaS has a lesser role in this condition. We also asked whether LtaS overexpression would restore LTA synthesis. The IPTG-inducible *ltaS* (*iltaS*) [9] established in the LAC strain (ANG2505; kindly provided by Dr. A. Gründling, Imperial College, UK) was grown overnight in SerFA containing 1 mM IPTG, without or with anti-FASII (AFN-1252, 0.5 µg/ml). Overnight cultures were then washed and restarted without or with both IPTG and AFN-1252; growth and LTA production were compared in the 4 conditions (**Fig. 4B**). Without IPTG, growth of the non-treated and anti-FASII-adapted *iltaS* strain nearly stopped, indicating that LtaS remained essential in both states. IPTG addition restored growth of non-treated and FASII^bypass^ cultures (**Fig. 4B**). However, IPTG-induced LtaS expression did not restore LTA production during FASII-bypass (**Fig. 4C**).

**Figure 4.**
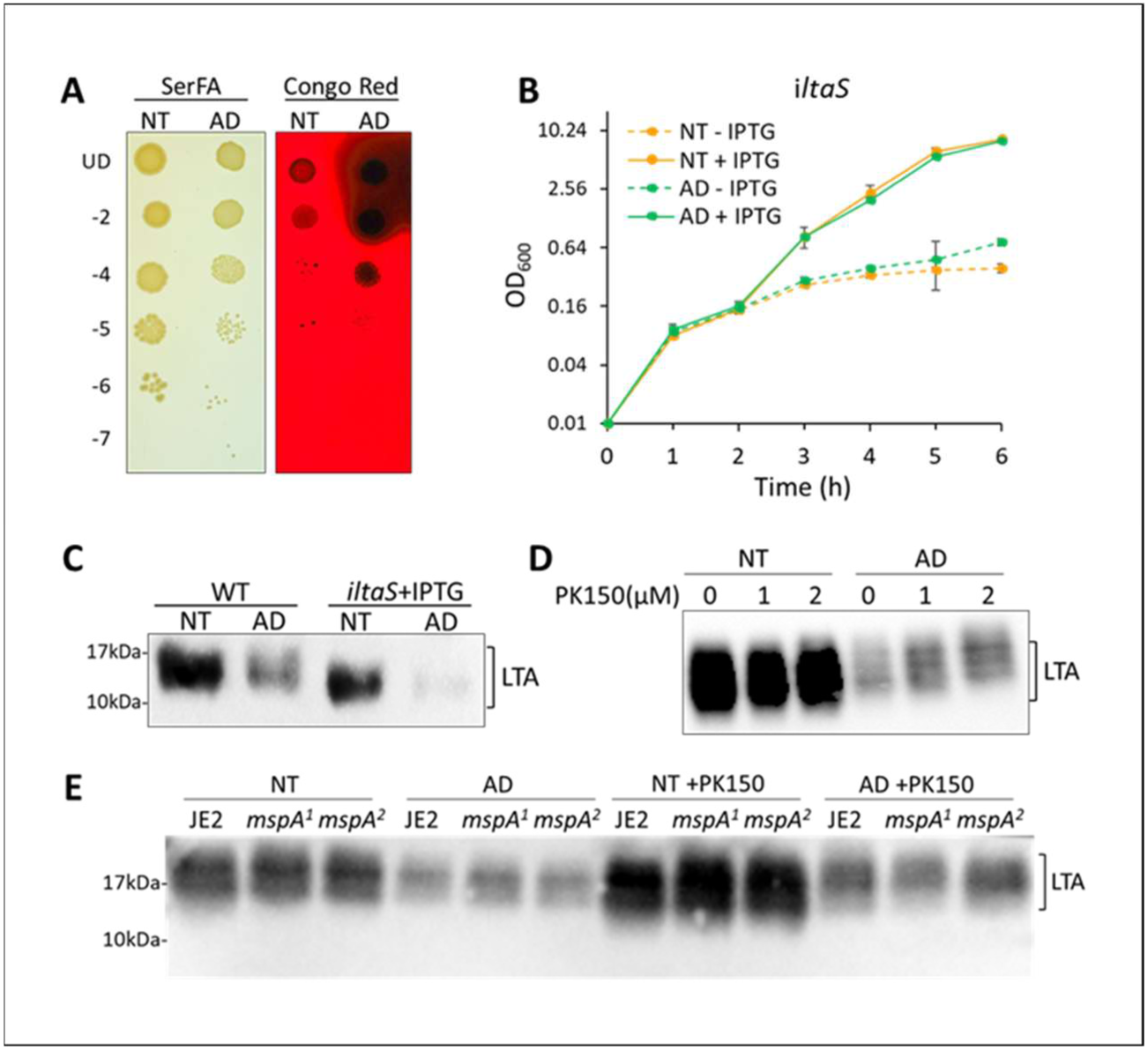
LtaS overproduction does not restore LTA during FASII^bypass^. **A.** Congo red inhibits LtaS activity [45]. Non-treated (NT) and anti-FASII-adapted (AD) *S. aureus* JE2 overnight cultures were adjusted to the same OD_600_ and dilutions were plated on solid SerFA medium without and with Congo Red (N=3). AD cultures show greater resistance than NT to Congo Red, possibly suggesting less reliance on LtaS. Black zones surrounding AD colonies on Congo Red indicates exopolysaccharide production [73, 74]. **B**, **C**. LtaS is required for growth but does not restore LTA in anti-FASII-adapted *S. aureus*. **B.** The IPTG inducible locus *iltaS* [9] established in the USA300 LAC derivative strain ANG2505 was grown overnight with 1 mM IPTG. Growth of ANG2505 NT (orange) and AD (treated by AFN-1252; green) cultures was followed without (dashed line) or with 1 mM IPTG (solid line). Results show the mean ± SD (standard deviation) of 4 biological replicates. **C**, **D**, **E**. LTA was detected by immunoblotting using anti-LTA antibody. **C**. Increasing LtaS expression does not restore LTA synthesis in FASII^bypass^ conditions. The WT parental LAC strain and ANG2505 *iltaS* were grown in NT and AD conditions in the presence of 1 mM IPTG. N=3. **D**. PK150 stimulates SpsB, which enhances LtaS cleavage and reportedly increases LTA [50, 51]. PK150 was added as indicated in NT and AD conditions and LTA production was assessed. **E**. Mutations in *mspA* derepress SpsB protease, leading to increased LTA [51]. LTA levels in JE2 WT and *mspA* mutants (USA300_FPR3757 mutant positions 2379899 [*mspA^1^*] or 2380097 [*mspA^2^*][63]) were examined in exponential phase SerFA (NT) or SerFA-AFN (AD) cultures. Samples at right were also treated with 1 µM PK150 during growth (N=3).

The SpsB protease cleaves LtaS and affects LTA length and localization [46–48]. The LtaS cleavage product does not synthesize LTA when alone [48], but the cleaved product appears to be required for efficient LTA synthesis [47]. We asked whether FASII^bypass^ creates conditions that reduce LtaS cleavage, which would explain LTA depletion. To test this, we used PK150 to stimulate SpsB [49, 50], and then assessed the effects on LTA synthesis. PK150 had no effect on LTA levels in either non-treated or FASII^bypass^ conditions (**Fig 4D**). MspA reportedly sequesters SpsB and *mspA* inactivation stimulates LTA production [51]. However, *mspA* mutants, without or with PK150, did not restore LTA during FASII^bypass^ (**Fig. 4E**). The above results show that neither LtaS nor LtaS maturation is limiting for LTA synthesis during FASII^bypass^. We therefore considered upstream pathways as possible causes for LTA depletion.

### FASII inactivation, but not membrane FA composition, dictates LTA depletion

FASII and/or FASII^bypass^ supplies FAs for phospholipid synthesis, which in turn provides the substrates for LTA synthesis (**Fig. 1**). We asked whether changes in FA composition during FASII^bypass^ might be responsible for LTA depletion. The *S. aureus* LTA lipid anchor comprises mainly two FAs, *ai*15 and C18:0, that also dominate membrane phospholipid composition [1]. In our initial studies showing LTA loss, medium was supplemented with a three-FA mixture (C14:0, C16:0, and C18:1), none of which are dominant in LTA. Possibly, these FAs cannot produce lipid anchors appropriate for LTA synthesis, which would explain decreased LTA production. We therefore performed the same experiment using a proportionately balanced mixture of FAs corresponding to those synthesized by *S. aureus* (C14:0, ai15, C16:0, C18:0, and C20:0; called ‘Natural FA mix’), so that the FA profiles of FASII^bypass^ and non-treated WT cultures were similar (**Fig 5A** upper). LTA levels were significantly decreased in both anti-FASII-adapted WT *S. aureus* and in the *fadD* deletion strain, even in the presence of this natural complement of FAs (**Fig. 5A**, lower). These results suggest that changes in FA composition are not a main factor leading to LTA depletion in FASII^bypass^ conditions.

**Figure 5.**
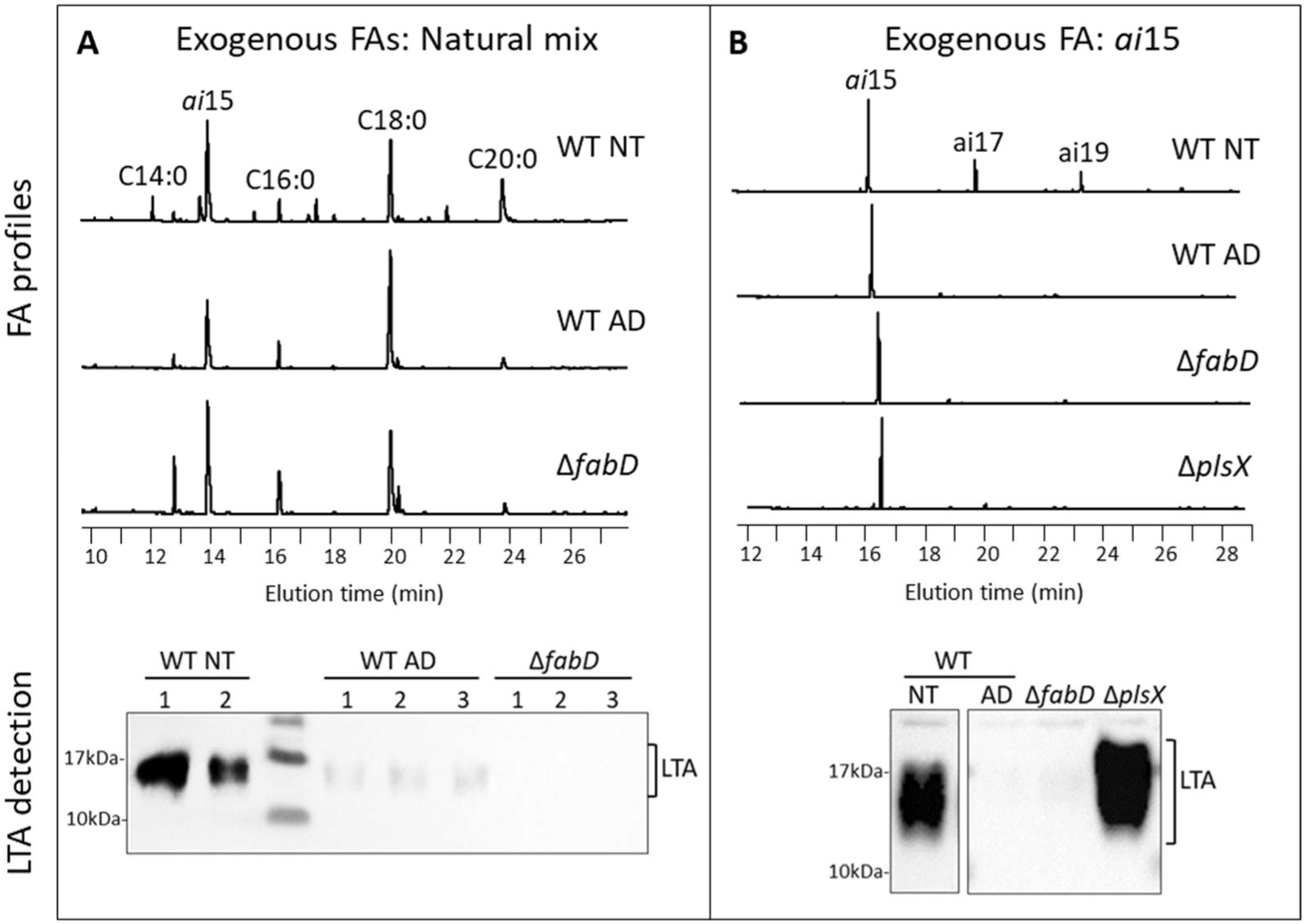
LTA production relies on FASII activity independently of membrane FA composition. *S. aureus* non-treated (NT), anti-FASII-adapted (AD), and Δ*fabD* (disabled for FASII) cultures were grown in **A**, 250 µM “Natural Mix”, which mimics endogenously produced FAs. **B.** The same strains and conditions as in **A**, plus a Δ*plsX* strain (deleted for *plsX*, see **Supplementary Fig. 1**; [52] were grown in 250 µM *ai*15, the major FA synthesized by *S. aureus*; *ai*15 is elongated to *ai*17 and *ai*19 only in the non-treated WT strain. **A**, **B**, upper: FA composition of membrane extracts determined by gas chromatography. Peak heights correspond to relative responses (mV) of each FA in a sample. Predominant FAs are indicated; N=2. **A**, **B**, lower: Cell extracts were prepared and submitted to immunoblotting using anti-LTA antibody N=3. Samples in **B** were run on the same gel and subjected to the same exposure time.

The above findings led us to hypothesize that FASII inactivation, and not the membrane FA composition, generates a condition that halts LTA production. To test this, we devised a means to obtain the same membrane FA composition in two conditions, one where FASII is inhibited (FASII^bypass^), and the other where FASII remains active. For the latter, we used a JE2-derived Δ*plsX* mutant devoid of PlsX [52, 53]. In Δ*plsX*, phospholipids comprise exogenous FAs in position 1 of the glycerophosphate backbone, and mainly FASII-synthesized *ai*15 in position 2. As *ai*15 is predominant in position 2, we obtained homogenous membrane FAs in FASII^bypass^ strains and in Δ*plsX* by supplementing growth media with solely *ai*15. Membrane FA profiles of Δ*plsX*, Δ*fabD*, and FASII^bypass^ cultures grown in *ai*15-supplemented media were nearly identical (**Fig. 5B upper**). In striking contrast, compared to both anti-FASII-adapted and Δ*fabD* cultures where LTA levels were depleted, LTA abundance in the Δ*plsX* mutant was similar to that in the non-treated WT strain (**Fig. 5B lower**). Therefore, changes in FA composition cannot account for LTA depletion. We conclude that inhibition of FASII activity, and not the membrane FA composition, is mainly responsible for LTA depletion.

### Pools of GroP, but not of ATP, are reduced during anti-FASII-adapted growth

The above results show that FASII activity is needed for complete LTA production, but not to provide the phospholipid precursors, which are assured by exogenous FAs. We asked whether the altered metabolite balance during FASII^bypass^ compared to FASII could account for the shift away from LTA synthesis. Examination of the major metabolites (**Fig. 6A**) involved in LTA production pointed to 2 candidates, ATP and GroP. ATP depletion leads to reduced LTA synthesis as shown in pioneering studies of Fischer on the biochemistry of LTA synthesis; synthesis of one LTA molecule costs ∼150 ATPs [1, 15]. However, ATP pools were higher in FASII^bypass^ compared to non-treated extracts, ruling out this explanation for LTA depletion (**Fig. 6B**). A neutral or positive ATP balance might be expected during FASII^bypass^, as incorporation of exogenous FAs economizes ATP cost of FA synthesis (estimated at ∼1 ATP per 2-carbon elongation [54]), e.g., 8 ATPs are used to synthesize one molecule of C16:0. This compares to just 1 ATP consumed by the Fak kinase [55] for incorporation of any length exogenous FA.

**Figure 6.**
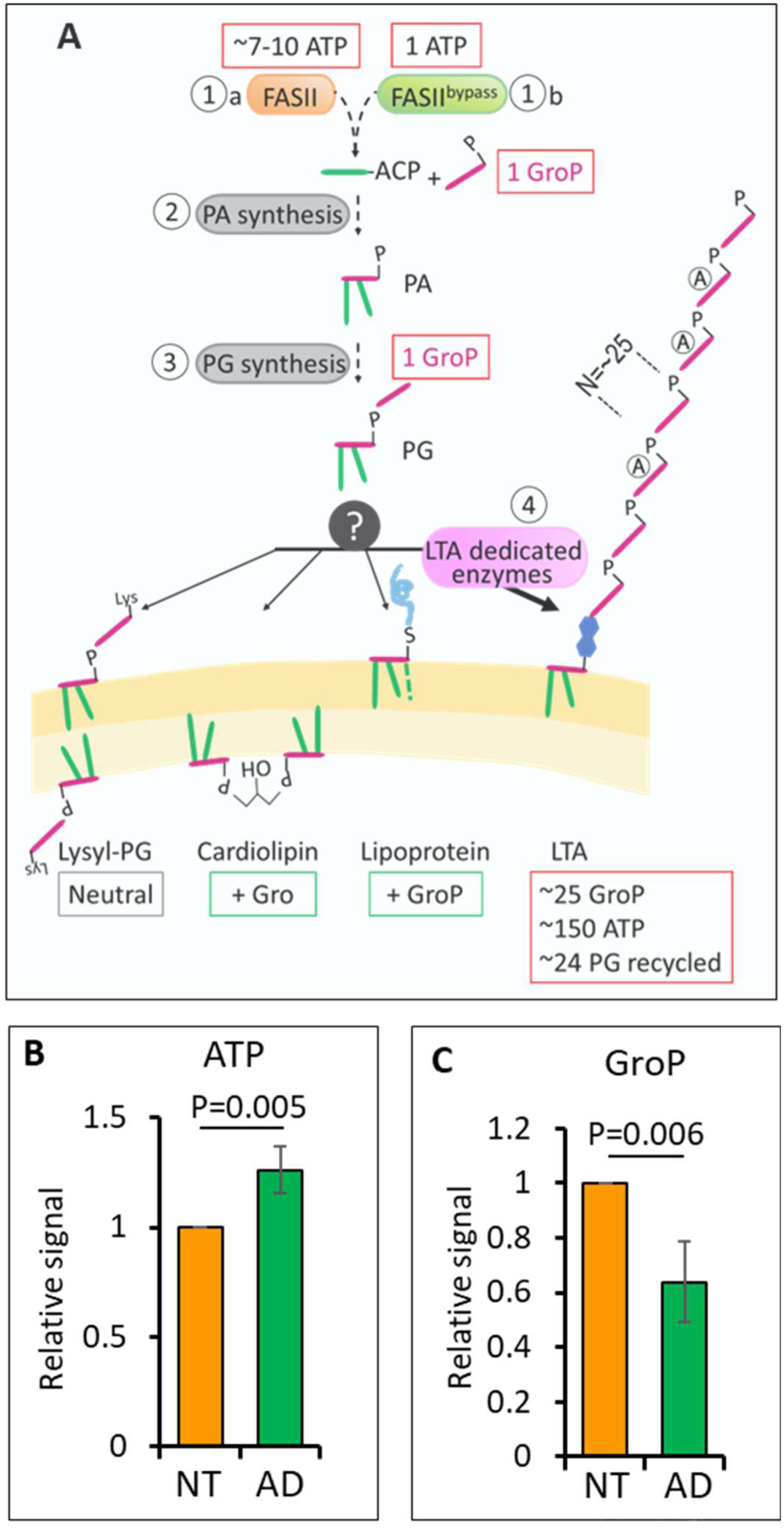
ATP and GroP costs of LTA synthesis, starting with FASII and FASII^bypass^ pathways. **A**. Multiple biosynthetic pathways underly synthesis of LTA, and carry an energy cost. LTA production relies on 3 major biosynthetic pathways (FASII and FASII^bypass^, ①a and ①b; phosphatic acid synthesis, ②; and phosphatidylglycerol synthesis, ③) before branching to LTA synthesis. FAs are in green; lipoprotein sometimes comprises a third FA (dashed line) [75]. ATP and GroP or Gro expenses (boxed in red), gains (in green), or neutral changes (in grey) during production of the 4 possible PG products, including LTA (Fig. 1). For a single LTA molecule, one PG is used to produce the anchor, while ∼25 PGs donate GroP to produce the polymer. The GroP donors are then recycled to regenerate PGs, which requires ATP. Synthesis of a single LTA molecule costs about 150 ATPs [1], 25 GroPs, and 26 PGs. In contrast, production of the three other lipid products leads to positive or neutral ATP and GroP footprints. PG product distribution in the inner and outer membrane leaflets is asymmetric: cardiolipin, but not the other lipid products, is preferentially enriched in the membrane inner leaflet [58]. **B** and **C**, bacteria were grown in SerFA (non-treated) or in SerFA-AFN (FASII^bypass^). **B.** ATP pools are greater in FASII^bypass^ conditions, as FASII^bypass^ uses ∼7-10 times less ATP per FA molecule compared to FASII (see **A**). N=5. **C**. FASII^bypass^ causes depletion of GroP pools. These measurements do not include GroP net loss associated with depletion of the GroP polymer attached to LTA. N=6. Data in **B** and **C** are shown as mean values ± SD normalized to cognate measurements in non-treated samples; P-values were determined using the two-sided T-test. Full data sets for **B** and **C** is in Fig. 6B and 6C Source data.

GroP is an essential substrate for both PG and LTA synthesis in *S. aureus*. Despite the low molar proportion of LTA compared to PG, about 50 % of total *S. aureus* GroP is sequestered within the LTA polymer in non-treated conditions [56]. Synthesis of one LTA GroP polymer involves sequential GroP transfer from ∼25 PG molecules (**Fig. 6A**; [1, 2]). As PG is the GroP donor, its synthesis necessarily precedes that of LTA. The GroP pool was about 36 % lower in the FASII^bypass^ condition than in non-treated bacteria (**Fig. 6C**). We conclude that the net drop in GroP, which is essential for PG synthesis followed by LTA GroP polymer synthesis, is the probable cause for LTA depletion in FASII^bypass^.

### Cardiolipin (CL) levels are increased in anti-FASII-adapted *S. aureus*, but do not affect LTA production

LTA is the only PG product to consume GroP (**Fig. 6A**). We asked whether other PG lipid products compensate LTA depletion in the GroP-limiting conditions imposed by FASII^bypass^. Whole cell lipids were extracted from non-treated, and FASII^bypass^ *S. aureus* cultures prepared in different FAs, i.e., ‘Natural FA mix’, *ai*15+C18:0 (the predominant FA moieties of LTA [1]), or *ai*15, the main *S. aureus* FA. The Δ*plsX* strain, which produces LTA, was also examined in the *ai*15 growth condition. The proportion of DG-DAG, the major LTA lipid anchor, decreased in FASII^bypass^ conditions, in keeping with both LTA and YpfP depletion (**Fig. 3**, **Supplementary Fig. S3**). In contrast, the proportions of CL increased to different extents according to the added FAs in all FASII^bypass^ *S. aureus* cultures (**Fig. 7A**). Of note, the lipid profile of Δ*plsX* was similar to that of the non-treated WT strain, despite comprising exclusively *ai*15 in its membrane (**Fig. 7A right**, violet bars); this indicates that FASII^bypass^ triggers these lipid changes.

**Figure 7.**
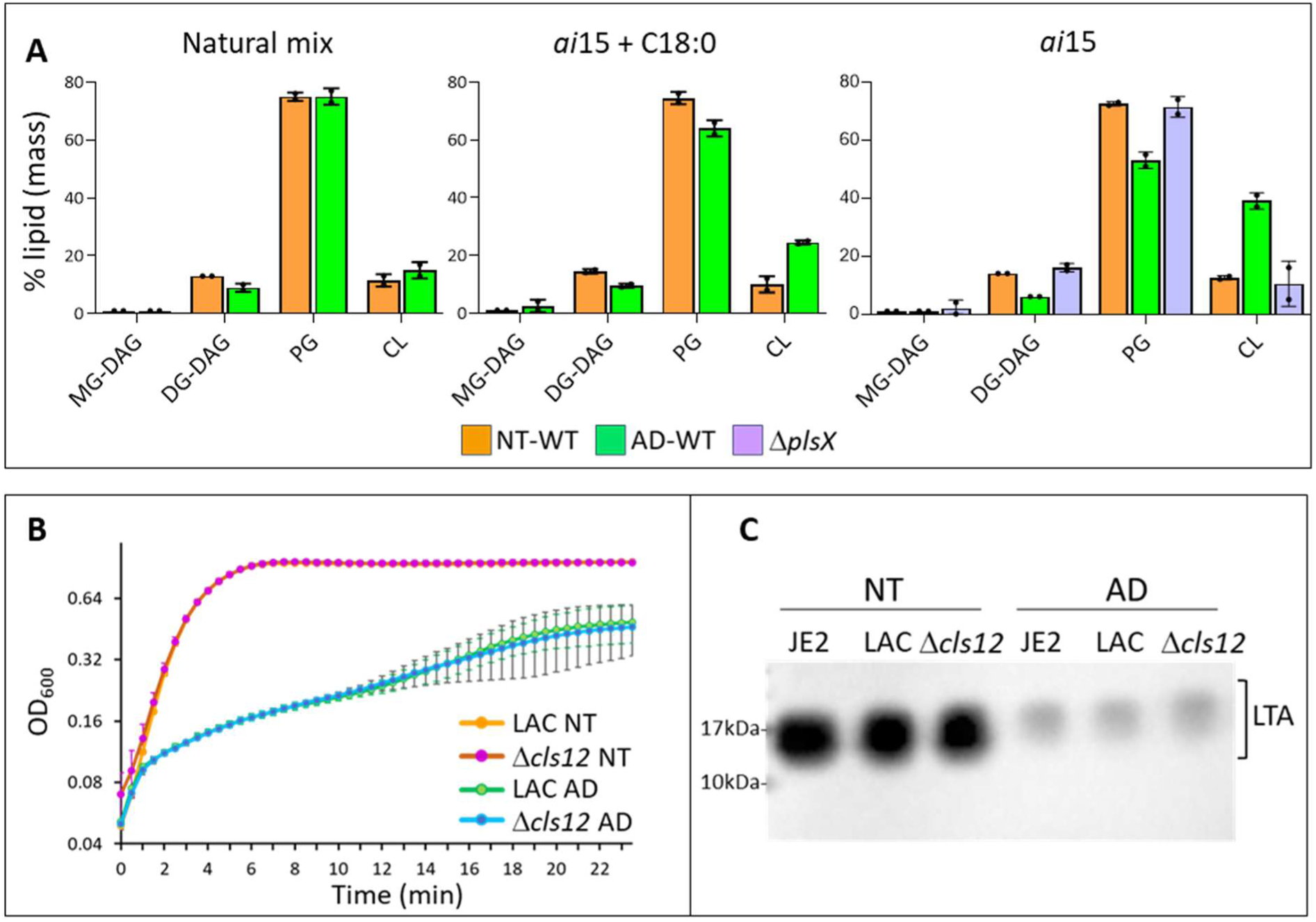
FASII^bypass^ leads to lipid species redistibution. **A**. Lipid extractions were performed on non-treated (NT) and anti-FASII-adapted (AD) *S. aureus* JE2, and the Δ*plsX* derivative where indicated, from OD_600_ = ∼3 cultures prepared in BHI plus 10% delipidated serum, supplemented by ‘Natural Mix’ (FA Mix), *ai*15 and C18:0 (125 µM each), or *ai*15 (250 µM). Relative mass proportions (% lipid (mass)) of monoglucosyl diacylglycerol (MG-DAG), diglucosyl diacylglycerol (DG-DAG; the LTA lipid anchor), phosphatidylglycerol (PG), and cardiolipin (CL) are shown, with data points (black dots), ranges, and average of biological duplicates. Of note, the Δ*plsX* strain produces LTA (Fig. 5) and its lipid distribution is similar to that of the WT NT strain. See Fig. 7A Source data for original data readouts. **B** and **C**. WT S. aureus JE2 and LAC strains, and the Δ*cls1*Δ*cls2* (Δ*cls12*) LAC derivative [57] were grown in SerFA without and with anti-FASII (AFN-1252; AFN). **B**. Growth of WT LAC and Δ*cls12* strains was compared in NT and AD cultures, and was comparable in each condition (N=3). **C**. LTA detection by immunoblotting was performed on WT JE2 and LAC, and Δ*cls12* cultures using anti-LTA antibody (N=3).

If CL were required during FASII^bypass^ as a compensatory mechanism, a cardiolipin synthesis mutant would fail to adapt to anti-FASII. We assessed FASII^bypass^ in a *cls1cls2* mutant, which lacks both cardiolipin synthases [57]. Growth and LTA depletion in FASII^bypass^ conditions were similar in WT and *cls1cls2* strains (**Fig. 7B** and **7C** respectively). Thus, while proportions of CL increase in response to anti-FASII, CL is not required for adaptation. This might be expected, as CL is enriched in the inner membrane leaflet (**Fig. 6A**; [58]), while LTA is exposed at the bacterial surface, ruling out a functional compensation. We therefore considered that non-lipid factors might support *S. aureus* growth during FASII^bypass^.

### WTA levels are increased in anti-FASII-adapted *S. aureus*, and required for adaptive growth and survival

Wall teichoic acid (WTA) and LTA are proposed to cooperate to assure bacterial envelope integrity, and WTA is required when LTA is absent [10, 16, 59]. Our previous proteomics heat map showed increases in 4 out of 5 detected WTA biosynthetic proteins in response to anti-FASII treatment (**Supplementary Fig. S3**). We assessed production of WTA as a possible factor compensating LTA loss during FASII bypass growth. *S. aureus* WTA and LTA yields were compared in non-treated and FASII^bypass^ conditions (**Fig. 8A** and **8B**, respectively). *S. aureus* FASII^bypass^ exhibited 2-fold greater WTA than the non-treated culture (p=0.05), while LTA amounts diminished by greater than 10-fold (p=5x10^-5^). Increased WTA production might contribute to *S. aureus* survival during anti-FASII treatment by compensating for reduced levels of LTA.

**Figure 8.**
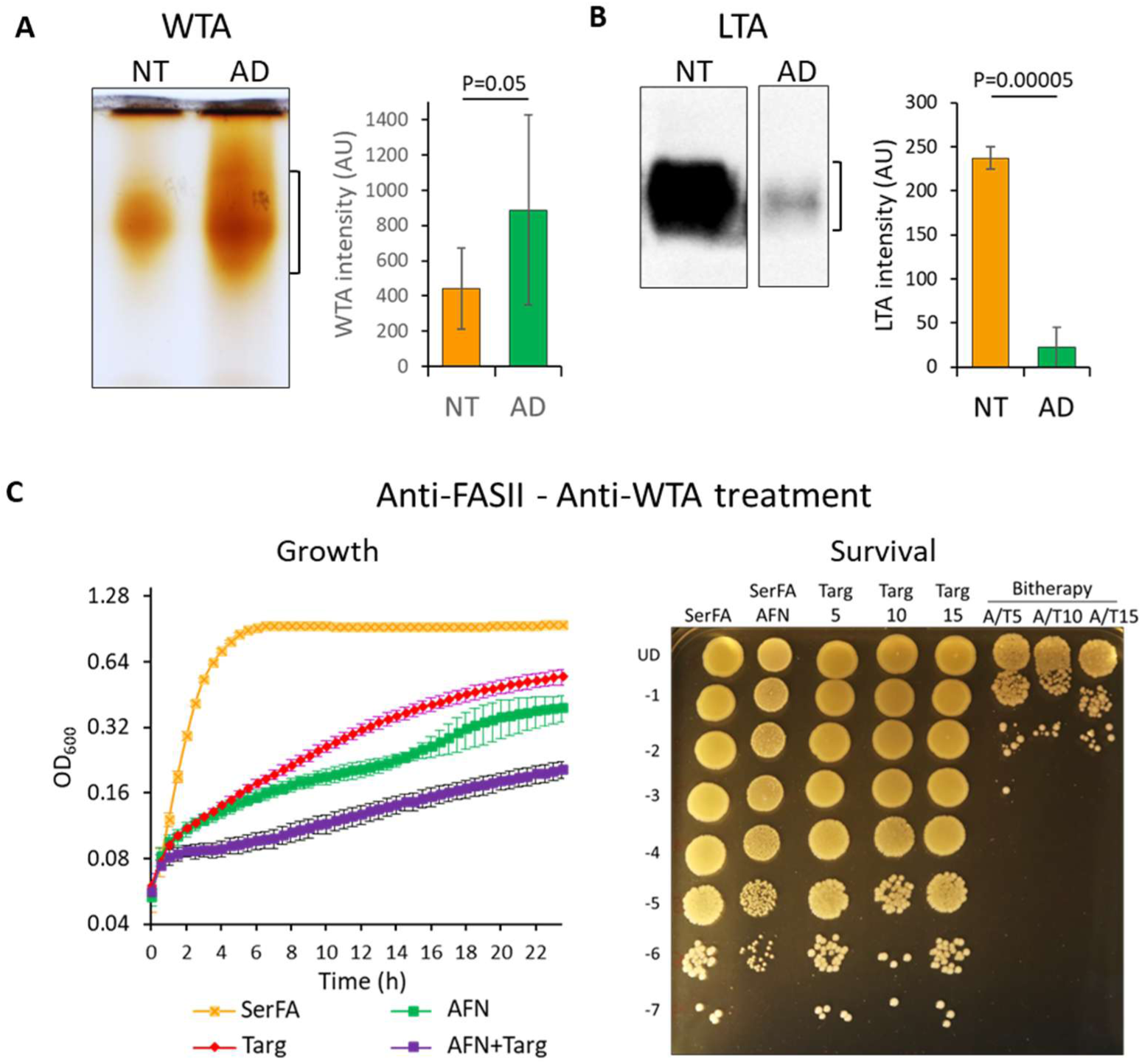
*S. aureus* wall teichoic acid (WTA) is produced at greater yields and is required for FASII^bypass^ growth and survival. **A** and **B**. Detection of WTA by Alcian blue staining (**A**) and LTA by immunodetection (**B**) in non-treated (NT) and anti-FASII-adapted (AD) *S. aureus* JE2 (N=6 and N=5 respectively). Representative gels, and means ± SDs of the independent samples are presented; P-values were determined using the two-sided T-test. **C**. Growth and survival of *S. aureus* upon bitherapy treatment with anti-FASII (AFN-1252, 0.5 µg/ml) and anti-WTA (Targocil, 15 µg/ml for growth, and 5-15 µg/ml for survival). Bacteria were precultured in SerFA prior to simultaneous antibiotic addition. Growth was monitored on a Tecan plate reader. Survival after 15 h growth was determined by serial dilution plating of cultures (5 µL per spot) that were treated with one or two antibiotics as indicated above each column.

As FASII-bypass reduces LTA and raises WTA levels, we asked whether FASII and WTA synthesis inhibitors would act synergistically. To test this, bitherapy assays were performed on *S. aureus* JE2, using AFN-1252 as anti-FASII, and targocil [60] as anti-WTA. Inhibition of growth and a >4-log reduction in 15 h survival indicate a synergistic biostatic effect of the combined treatment (**Fig. 8C**). These promising results suggest a bitherapy strategy to enhance the effects of two drugs that reach their targets, where single treatment is ineffective.

## Discussion

The FA building blocks of all membrane phospholipids are synthesized by FASII and/or obtained from environmental sources via FASII^bypass^. We showed that while both pathways generate FAs and phospholipids, FASII, but not FASII^bypass^, promotes LTA production. The FASII activity requirement for LTA production elucidated here in *S. aureus* also applies to a streptococcus (*S. agalactiae*), where external FAs repress FASII [32], suggesting the generality of FASII-controlled LTA synthesis.

Two explanations for the differential effects of FASII and FASII^bypass^ on LTA synthesis were excluded. First, a regulatory effect of FASII on the essential LTA synthesis enzyme LtaS was ruled out, as LtaS overproduction did not restore LTA during FASII^bypass^ (**Fig. 4**). Second, a change in membrane FA composition during FASII^bypass^ did not cause LTA depletion, as the same FA composition obtained by manipulating two different pathways, one being FASII^bypass^, and the other conserving FASII but inactivating PlsX, resulted in different LTA outcomes (**Fig. 5B**). These exclusions led us to focus on metabolites potentially underlying LTA synthesis that might be dictated by FASII and FASII^bypass^ pathways.

PG, the major *S. aureus* phospholipid, is the substrate for four possible products (**Fig. 1**). Among them, LTA synthesis is doubly costly: First, LTA hordes an estimated half of bacterial GroP in polymer form, which it receives from ∼25 PGs. Second, by transferring their GroP moieties, the PGs must be recharged with GroP in an ATP-dependent process [1, 56, 61]. LTA is the only PG product that calls for this costly tradeoff, a case of “deshabiller Pierre pour habiller Paul”, where numerous PGs are dismantled to produce one LTA. Our findings showing GroP depletion by FASII^bypass^ lead to a simple model in which the ‘choice’ of synthesizing LTA depends on the amount of available GroP, and on whether PG production is slowed due to insufficient GroP (**Fig. 6A** and **6C**). Reduced LTA synthesis could favor production of competing PG products that do not use GroP, i.e., CL, lysyl-PG, and/or lipoprotein. Higher proportions of CL detected during FASII^bypass^ are consistent with this proposal. These results connect the bacterial metabolic state to expression levels of a multifunctional surface structure, LTA, which contributes to cell integrity and division, and virulence. They tie in with our previous observations showing slowed virulence kinetics of FASII^bypass^ adapted bacteria [35].

The reasons for the reduction in GroP pools upon FASII^bypass^ are not yet known. Unlike FASII, FASII^bypass^ relies on reverse PlsX activity (**Fig. S1**), which alters enzyme-product homeostasis, and causes accumulation of phospholipid intermediates [33, 34]. Previous work in *Bacillus subtilis* showed that accumulated lysophosphatidylglycerol (LPA), a phospholipid intermediate, is unstable, generating dephosphorylated monoacylglycerol; the GroP moiety is lost in this process [62]. We speculate that FASII^bypass^ and consequent PlsX reversal desynchronizes phospholipid synthesis enzymes, similarly leading to LPA accumulation and dephosphorylation of the GroP moiety. Futile GroP turnover from phosphatidic acid (PA) intermediates would explain its depletion during FASII^bypass^ (schematized in **Supplementary Fig. S6**, salmon color inset).

Rapid growth of *S. aureus* and *S. agalactiae* by FASII^bypass^ indicates that LTA depletion is compensated by other factors, as also suggested from the numerous suppressor mutants and conditions that alleviate the LTA requirement [25, 27–31]. CL is increased, but not required for FASII^bypass^; this might be expected, due to its enriched location in the inner membrane leaflet [58], rather than the outer leaflet where LTA is located. The roles of other outer leaflet lipids, i.e., lipoproteins, in compensating LTA depletion remain to be explored. The overall increase in WTA yields and accrued sensitivity to a WTA synthesis inhibitor point to a more dominant role for WTA during FASII^bypass^, likely by compensating LTA depletion (**Fig. 8**). Simultaneous inactivation of LTA and WTA is lethal, possibly due to a collapse of the protective stiff cell wall structure that requires at least one of these components [16, 18, 59]. Our findings showing the collateral effects of anti-FASII in altering *S. aureus* virulence factor expression and reducing LTA can be exploited to design synergistic antimicrobial bitherapy, combining an anti-FASII and an anti-WTA, to eliminate *S. aureus*.

## Materials and Methods

### Strains, media, and growth conditions

The following *S. aureus* strains were used: USA300_FPR3757 JE2 [63]); RN-R and HG1-R, which are derivatives of RN4220 and HG001 respectively, whose *fakB1* alleles were repaired; note that the NCTC 8325 lineage carries a deletion in *fakB1*, resulting in a defect in exogenous FA utilization [64]. A USA300 Δ*fabD* mutant (removing the malonyl CoA-ACP transacylase) was created as described [40]. The following strains were generously provided as follows: strain ANG2505, IPTG-inducible expression of *ltaS*, described in RN4220 [9], and established in USA300 LAC, from Dr. Angelika Grundling (Imperial College London, UK); *mspA* transposon insertion mutants corresponding to USA300_FPR3757 positions 2379899 or 2380097, from Dr. Paul Fey (University of Nebraska, USA) [63]; USA300_FPR3757-derived LAC strain with a double deletion of *cls1* and *cls2*, from Dr. Andreas Peschel (University of Tubingen, Germany) [57]. ANG2505 and *mspA* mutants were selected on 5 µg/ml erythromycin prior to experiments. Strains were cultured in BHI, or SerFA (BHI comprising 10 % decomplemented calf serum with 250 µM FAs (Laradon, Sweden) prepared as an equimolar mixture of C14:0, C16:0, and C18:1). Where specified, single FAs or a mixture simulating the FAs endogenously produced by *S. aureus* (‘natural mix’ containing C14:0, 6.5%; ai15, 40.4%; C16:0, 6.3%; C18:0, 34.1%; C20:0, 12.6%) were prepared in BHI containing 10% delipidated bovine serum (Eurobio, France) and used at the 250 µM final concentration. Cultures were started from independent colonies from BHI solid medium, and inoculated overnight as SerFA pre-cultures grown aerobically at 37°C in SerFA. They were then diluted in SerFA to OD_600_ = 0.1 without or with antibiotics, and growth was followed for at least 10 h. FASII inhibitors were AFN-1252 (anti-FabI; Bioaustralis, Australia) or platensimycin (anti-FabF, MedChemExpress, France), both used at 0.5 µg/ml. PK150 (Tebubio, France), which stimulates SpsB activity [50], was used at 1-2 µM as indicated. *S. aureus* growth experiments were performed with aeration at 37°C in tubes or using a plate reader (Tecan Spark, Tecan France). For the latter, cultures were deposited in 96 well plates at initial OD_600_ = 0.05 and growth was monitored for 24h at 37°C. All growth experiments were performed in at least three biological replicates. *S. agalactiae* NEM316 [32] was grown at 37°C without shaking in BHI containing 0.25% BSA without or with 100 µM C17:1. This FA represses FASII without deterring growth [43, 65].

### Electron microscopy

Bacteria were fixed with 2% glutaraldehyde in 0.1 M Na cacodylate buffer (pH 7.2) for 3 hours at room temperature. Samples were then contrasted with 0.2% Oolong Tea Extract in cacodylate buffer, postfixed with 1% osmium tetroxide containing 1.5% potassium cyanoferrate, and gradually dehydrated in ethanol (30% to 100%). The samples were substituted in a mixture of ethanol and Epon and embedded in Epon resin. Thin sections (70 nm) were collected onto 200 mesh copper grids and counterstained with lead citrate. The grids were examined using a Hitachi HT7700 electron microscope at 80 kV, and images were captured using a charge-coupled device camera. Envelope thickness was assessed on 25-30 cells and at least 2 measurements were done per cell using ImageJ FIJI software. and calculations of mean, standard deviation, and 2-tailed T-tests used for statistical significance were performed using GraphPad Prism 9.5.1 and Excel software.

### Ethidium bromide retention

*S. aureus* JE2 was precultured in SerFA and transferred to fresh medium without or with AFN-1252, to obtain exponential (OD_600_ = 1) and stationary (overnight) cultures. 0.5 OD_600_ units were collected for each sample and washed twice in PBS in the same volumes. 100 µl of each washed sample was transferred to a black 96-well plate, and ethidium bromide (1 µg/ml) was added. Ethidium bromide retention was monitored essentially as described [66, 67] on a Tecan plate reader at 37°C for 60 min, using excitation wavelength 539 nm and emission wavelength 600 nm. The 20 min time point was used to assess differences between samples. Calculations of mean, standard deviation, and 2-tailed T-tests used for statistical significance were performed using Excel software.

### LTA Detection by immunoblotting

LTA in *S. aureus* USA300 JE2 and LAC strains, and *iltaS* was assessed in the specified OD_600_ and conditions. Cultures were inoculated in SerFA and then subcultured overnight in SerFA without or with anti-FASII. The following day, cultures were transferred to the same fresh media and grown to OD_600_ between 1 and 3. Ten OD_600_ units of each culture were collected and centrifuged at 8,000 rpm for 5 min, washed once with phosphate-buffered saline (PBS) containing 0.02% Triton, and then washed twice with PBS. Pellets were stored at -80 °C prior to cell extractions. FastPrep (program 2; 6.5 m/s, 50 s x 3) was applied to all samples. Bradford protein assay was performed on lysates to determine sample concentrations. Ten µg equivalents of protein were deposited in wells of 15% polyacrylamide-SDS gels. Gel contents were transferred to a PVDF membrane (BIO-RAD, France) using the Power Blotter System cassette (Thermo Scientific, France) with the ‘medium’ molecular weight program for 7 minutes. PVDF membranes were incubated with primary LTA-specific antibody (Clone 55; HyCult Biotechnology, Holland) and horseradish peroxidase-conjugated goat anti-mouse secondary IgG antibody (ThermoFisher, France), diluted to 1:2,500 and 1:10,000, respectively. Immunoreactive LTA species were detected using enhanced chemiluminescence (ThermoScientific, France), analyzed with the Image Lab 5.0 software (ChemiDoc MP Imaging System, BioRad, France), and quantified with ImageJ Fiji software [68]. LTA immunodetection experiments were performed on independently prepared samples at least 10 times in WT JE2 in both non-treated and FASII^bypass^ conditions, and as indicated for other *S. aureus* and *S. agalactiae* assays.

### Congo Red resistance

Non-treated and anti-FASII-adapted *S. aureus* JE2 cultures were grown to OD_600_ ∼2 and then adjusted to OD_600_ = 1 to perform spot test dilutions (undiluted=UD). For each dilution, 5 µl aliquots were spotted onto SerFA solid medium without or with 2.5 mg/ml Congo Red. Plates were incubated 24 h at 37°C, and photographed.

### FA Extraction

The equivalent of one OD_600_ unit of bacterial culture was centrifuged at 8,000 rpm for 5 min, washed once in 0.9% NaCl containing 0.02% Triton X-100, then twice in 0.9% NaCl at the same speed and time. Cell pellets were subjected to membrane lipid extraction as described [33, 69]. Gas chromatography was performed in split-splitless injection mode on an AutoSystem XL Gas Chromatograph (Perkin-Elmer) equipped with a ZB-Wax capillary column (30 m x 0.25 mm x 0.25 mm; Phenomenex, France). Data were analyzed using the TotalChrom Workstation program (Perkin-Elmer). *S. aureus* and *S. agalactiae* FA peaks were detected between 12 and 32 minutes of elution and identified based on their retention times compared to purified esterified FA standards.

### Lipid Extraction and Profiling

For lipid extraction, 100 OD_600_ unit equivalents were collected from bacteria grown to OD_600_ ≈ 3. Bacteria were washed as above for FA extraction. Pellets were freeze-dried overnight and stored at -80°C. Lipid extractions were performed as described [70]; all steps were performed in glass tubes. Briefly, pellet lipids were extracted with 4.75 ml of extraction buffer (chloroform, methanol, and 0.3 % NaCl in a 1:2:0.8 ratio) incubated at 80°C for 15 min, followed by 1 h vortexing at room temperature. This procedure was repeated once, but with a 30 min vortexing step. Then, 2.5 ml each of chloroform and 0.3 % NaCl were added consecutively, and phase separation was achieved by 15 min centrifugation at 4,000 rpm. The lower phase was collected and evaporated under nitrogen gas. Dried total lipid extract was weighed and stored at -20°C. Prior to lipid analysis, samples were solubilized in chloroform to obtain a concentration of 70 mg/mL of dried extract.

Lipid analysis was performed using Normal Phase Liquid Chromatography (NPLC) as described [71] with a Dionex Ultimate 3000 RSLC system (ThermoFisher Scientific, Germany) equipped with two quaternary pumps, an autosampler, and a column oven. The RSLC system was coupled online to a Corona Ultra charged aerosol detector (Corona-CAD) and to an LTQ Orbitrap Velos Pro mass spectrometer equipped with a linear ion trap and an orbital trap analyzer (ThermoFisher Scientific, Germany). Lipid classes were quantified by using a mixture of commercial standards (Avanti, Germany) in chloroform, containing mono-glucosyl diacylglycerol (MG-DAG, 840523P), di-glucosyl diacylglycerol (DG-DAG, 840524P), phosphatidylglycerol (PG, 841138P) and cardiolipins (CL, 840012P); the equimass mixture was injected at concentrations ranging from 0.025 to 0.5 mg/mL for each component. Cyanurphosphatidylethanolamine (Cyanure-PE(16:0-16:0), 870287P, Avanti, Fr) was used as an internal standard. Five µL samples were injected and concentrations of each lipid class were determined with the calibration curves obtained by corona-CAD detection. Lysyl-phosphatidylglycerol (Lysyl-PG) was quantified using the PG calibration curve. Mass spectrometry was used to confirm the lipid species detected and estimate its average molecular mass. Data were analyzed using Thermo Xcalibur Qual Browser. The proportion of each lipid is presented as the mean ± SD of two biological replicates using GraphPad Prism 9.5.1.

### ATP measurements

Non-treated and anti-FASII-adapted cultures were grown in SerFA medium to OD_600_ = 2. For each sample tested, 100 µl of culture was collected and placed in a 96 white opaque-multiwell plate (Nucleon, France). 100µl of Bactiter-Glo (Promega, France) was added to each sample. The plate was incubated at 25°C with shaking (160 rpm) for 5 min before measuring luminescence (Tecan plate reader). Calculations of relative expression, standard deviation, and 2-tailed paired T-tests used for statistical significance were performed using Excel software.

### GroP measurements

JE2 cultures were prepared in non-treated (NT) and anti-FASII-adapted (AD) conditions in SerFA medium, supplemented with AFN-1252 (0.5 µg/ml) for AD strains. The following day, cultures were diluted to OD₆₀₀ = 0.05 in fresh medium and grown to OD₆₀₀ = ∼3. Twenty-five OD_600_ equivalent units were collected and centrifuged at 8000 rpm for 5 min at room temperature. Pellets were flash-frozen in liquid nitrogen to halt enzymatic reactions. Cells were then treated with 1 mL 5% perchloric acid (HClO₄) for 30 minutes on ice, followed by thorough resuspension. Lysates were centrifuged at 12100 rpm (14000 g) for 30 minutes at 4°C, and supernatants were collected and stored at -80°C until analysis.

GroP was quantified by liquid chromatography coupled with mass spectrometer detector (LC-MS). Samples were injected for liquid chromatography using a Supelco-F5 column (2.1 × 150 mm; 3 µm) with a mobile phase consisting of water with 0.1% formic acid (phase A) and acetonitrile with 0.1% formic acid (phase B). Elution was performed using a gradient program as follows: the run starts at 5% phase B for 2 minutes, then increases to 100% phase B over 13 minutes, maintaining these conditions for 6 minutes before returning to 5% phase B in 1 minute, followed by re-equilibration for 10 minutes. The flow rate was set at 0.25 mL/min, and detection was carried out by mass spectrometry using an Nx8060 triple quadrupole ion analyzer (Shimadzu, France) equipped with an electrospray ionization (ESI) probe operating at 250°C. Detection was performed in multiple reaction monitoring (MRM) mode, with the following transitions: 171.15>79.00; 171.15>127.10 and 171.15>103.05 (in negative mode) and 173.10>99.05;173.10>81.10 and 173.10>63.05 (in positive mode). Calibration curves were previously generated using commercial standards in the range of 0.2 to 50 pmol injected. Calculations of relative expression, standard deviation, and 2-tailed paired T-tests used for statistical significance were performed using Excel software.

### WTA detection

Non-treated and anti-FASII-adapted *S. aureus* JE2 were prepared in SerFA without or with AFN-1252 (0.5µg/ml) and cultured to OD_600_ = 2-3. Extractions were performed on 60 or 180 OD_600_ equivalent units of bacterial cultures as described [72]. Samples were migrated on 15% native polyacrylamide gel electrophoresis (PAGE) and visualized using Alcian blue and silver nitrate staining as described [72]. Gels were scanned on an Epson scanner and WTA peaks were quantified using ImageJ Fiji software [68]. Standard deviation, and 2-tailed paired T-tests for statistical significance were performed using Excel software.

### Combination anti-FASII and anti-WTA treatment effects on *S. aureus* growth and survival

Growth of JE2 was assessed in SerFA supplemented or not with AFN-1252 (0.5µg/ml) without or with the TarG inhibitor targocil (anti-WTA, 15 µg/ml) [60]. Cultures were deposited in 96 well plates at initial OD_600_ = 0.05 and growth was monitored for 24 h at 37°C using a Tecan plate reader. Results correspond to means of three biological replicates. Survival to single and combined treatment was performed using the above growth conditions, except that targocil was added at 0, 5, 10 or 15 µg/ml. Bacterial survival was assessed by serial dilution platings after 15h growth. All experiments were performed in triplicate.

## Supporting information

Source file for Fig. 2A

Source file for Fig. 2B

Source file for Fig. 6B and 6C

Source file for Fig. 7A

## Acknowledgements

We are grateful to Angelika Gründling (Imperial College London, UK) for insightful discussion of this work. We thank Professors A. Gründling, Andreas Peschel (University of Tubingen, Germany), and Paul Fey and Jennifer Endres (University of Nebraska) for their generous gifts of strains. We thank Marie-Françoise Noirot-Gros and Hasna Toukabri (Micalis), and team colleagues Jasmina Vidic, Philippe Gaudu and David Halpern for valuable discussions and advice concerning this work. PW was awarded a Franco-Thai PhD scholarship from Campus France. We gratefully acknowledge funding from Fondation pour la Recherche Medicale (DBF20161136769; AG), the Agence Nationale de la Recherche (ANR-16-CE15-0013; AG, PTC), and the ANR under the umbrella of the Joint Programming Initiative on Antimicrobial Resistance (JPIAMR) ANR funding (ANR-22-AAMR-0007; AG, PTC). We thank the Région Ile-de-France for financial support in the acquisition of instruments for the SAMM core facility (AS, BP).

## Supplementary Figure Legends

**Supplementary Figure S1. *S. aureus* has two ways to produce FAs for phospholipid synthesis: FASII and FASII^bypass^. A**. Both FASII and FASII^bypass^ provide FAs for phospholipid synthesis, but with different outcomes. The FASII pathway is energetically costly, and provides acyl-acyl carrier protein (FA-ACP) for phospholipid synthesis. FASII^bypass^ incorporates environmental FAs (eFA) that produce FA-ACP *via* reverse PlsX activity. LPA, lysophosphatidic acid; PA, phosphatidic acid; GroP is represented by a burgundy line joined to a P representing the phosphate group. **B**. Growth of *S. aureus via* FASII in SerFA (non-treated, orange), or upon exposure to a FASII inhibitor (AFN-1252 0.5 µg/ml, green curve), when it incorporates exogenous FAs. An initial lag period is followed by robust and sustained growth in a non-mutational response [33, 35]. Growth curves are representative of >20 determinations. Refers to information from Introduction.

**Supplementary Figure S2. Transmission electron microscopy of *S. aureus* JE2 in non-treated and anti-FASII-adapted growth at 6 and 10 hours.** Bar, 200 nm. Micrographs complement those shown in Fig. 2.

**Supplementary Figure S3. Kinetic heatmap of *S. aureus* envelope synthesis proteins whose levels are altered during FASII^bypass^.** Proteomics analyses were performed previously on *S. aureus* USA300 JE2 strain grown in SerFA, and treated or not with the FabI inhibitor triclosan (0.5 µg/ml), performed on biological quadruplicates (from [35]). Sampling times correspond to 2, 4, 6, 8, and 10 h post anti-FASII-treatment. The heat map shows changes in detected LTA and WTA biosynthetic enzymes, and D-alanylation enzymes, which mediate decoration of both structures [76]. Changes in protein expression were determined relative to weighted values for each protein (scale at left: navy, down-represented and yellow, up-represented). WTA biosynthetic enzymes TarL’ (also called TarK) and TarL are redundant [77]. Data is derived from [35].

**Supplementary Figure S4. LTA remains depleted after prolonged incubation in anti-FASII-adapted *S. aureus*. A**. *S. aureus* JE2 was grown in SerFA medium without and with AFN-1252 and samples were harvested at 6 or 10 hours, or after overnight (ON) growth. The ON anti-FASII-adapted cultures were sub-cultured (Subc 1) into fresh SerFA-AFN or SerFA medium (indicated by curved arrows) and grown 3 h. A second subculture (Subc1 to Subc 2) was prepared in the same condition. Whole cell extracts were prepared and LTA was detected by immunoblotting using anti-LTA antibody. LTA remains depleted even after long term adaptation (N=3). **B**. Samples from **A** were extracted for FA analyses at the indicated OD_600_ or time. Red, exogenous FAs; black, major endogenous FAs (N=2). Supports data from Fig. 3.

**Supplementary Figure S5. LTA detection in *S. aureus* NCTC 8325 derivatives**. *S. aureus* RN4220 and HG001 are both from the NCTC 8325 lineage, which lacks the functional *fakB1* gene required for complete exogenous FA incorporation [78]. Repair of *fakB1* generated RN-R and HG1-R respectively [64]. Cultures were grown in SerFA without (non-treated, NT) and with AFN-1252 (anti-FASII-adapted, AD). LTA production was detected by immunoblotting using anti-LTA antibody (N=3). HG1-R samples are from a single gel, immunoblot, and exposure time. Supports data from Fig. 3.

**Supplementary Figure S6. Model linking FASII synthesis arrest to GroP metabolite depletion.** FASII inhibition due to antibiotics, mutation, and/or exogenous FA inhibition, is compensated by incorporation of exogenous FAs in membranes, schematized here for *S. aureus* [32–34, 43, 65]. Incorporation requires FakAB for FA phosphorylation [79], and reverse PlsX activity to provide acyl-ACP [33, 34]. This rerouted cycle is proposed here to desynchronize acyl-ACP and lysophosphatidic acid (LPA) availability for PlsC-mediated PA synthesis. Accumulated LPA intermediates are unstable, leading to degradation of their Gro-P moieties (salmon color zone; [62]), consistent with Gro-P depletion in anti-FASII-adapted S. aureus (Fig. 6). We propose that FASII blockage leads to GroP breakdown, such that the high demand for GroP from PG turnover to produce LTA cannot be met. Short arrows indicate the coordinate LTA decrease and cardiolipin increase when FASII is inhibited. Strong black arrows, favored reactions; thin black arrows, reduced or inhibited reactions; dashed arrow, multi-step reactions. Red arrows and pink zone highlight reactions leading to Gro-P depletion. The upper part of the figure applies information from [62]. The model supports results from Fig. 6.

Source data is available for results from Fig. 2A, 2B, Fig. 6B, **6C**, and Fig. 7A.

