## Supplementary Figs 1-6 for "The fatty acid synthesis pathway is a checkpoint for lipoteichoic acid synthesis in *Staphylococcus aureus*"

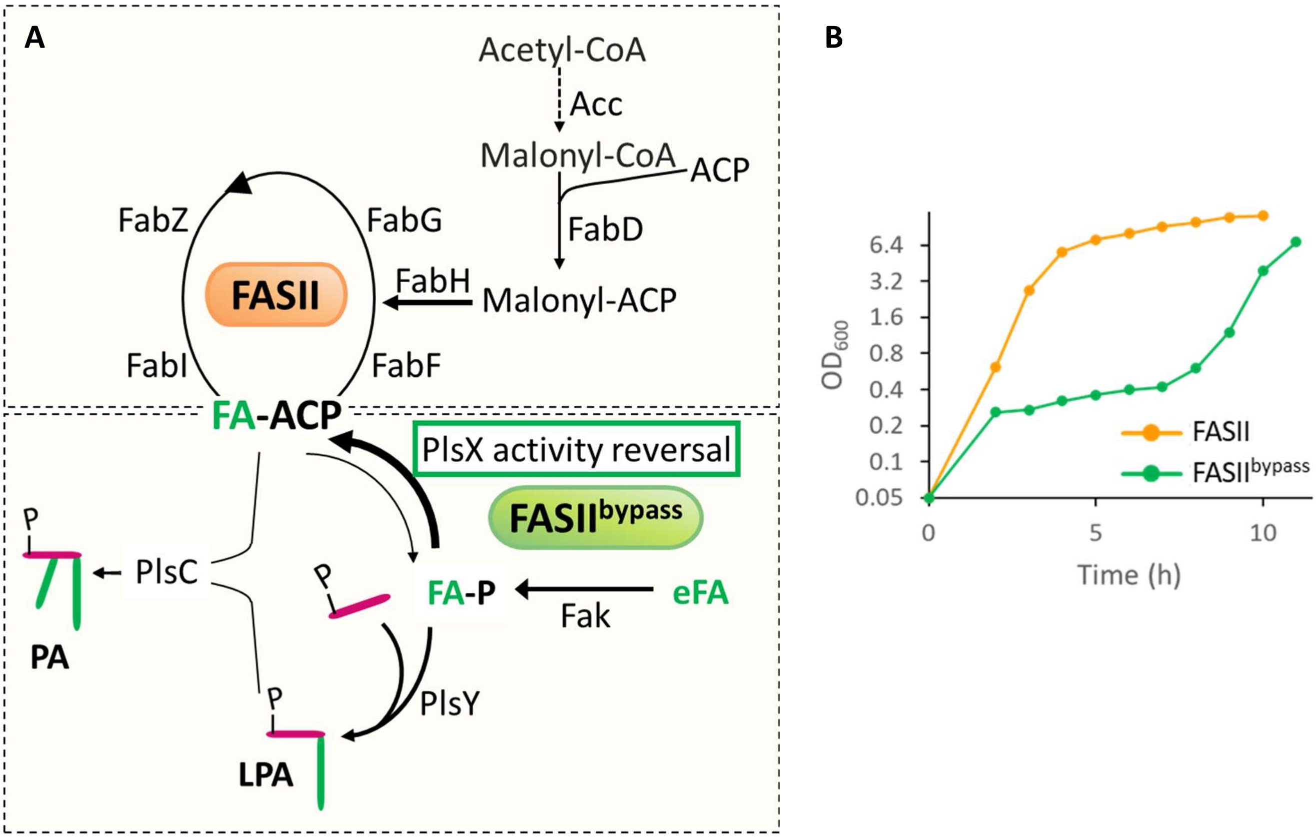


**Supplementary Fig. S1**. ***S. aureus*** **has two ways to produce FAs for phospholipid synthesis: FASII and FASII$^{bypass}$. A.** Both FASII and FASIIbypass provide FAs for phospholipid synthesis, but with different outcomes. The FASII pathway is energetically costly, and provides acyl-acyl carrier protein (FA-ACP) for phospholipid synthesis. FASIIbypass incorporates environmental FAs (eFA) that produce FA-ACP via reverse PlsX activity. LPA, lysophosphatidic acid; PA, phosphatidic acid; GroP is represented by a burgundy line joined to a P representing the phosphate group. **B.** Growth of *S. aureus* via FASII in SerFA (non-treated, orange), or upon exposure to a FASII inhibitor (AFN-1252 0.5 µg/ml , green curve), when it incorporates exogenous FAs. An initial lag period is followed by robust and sustained growth in a non-mutational response [33, 35]. Growth curves are representative of >20 determinations. Refers to information from Introduction.

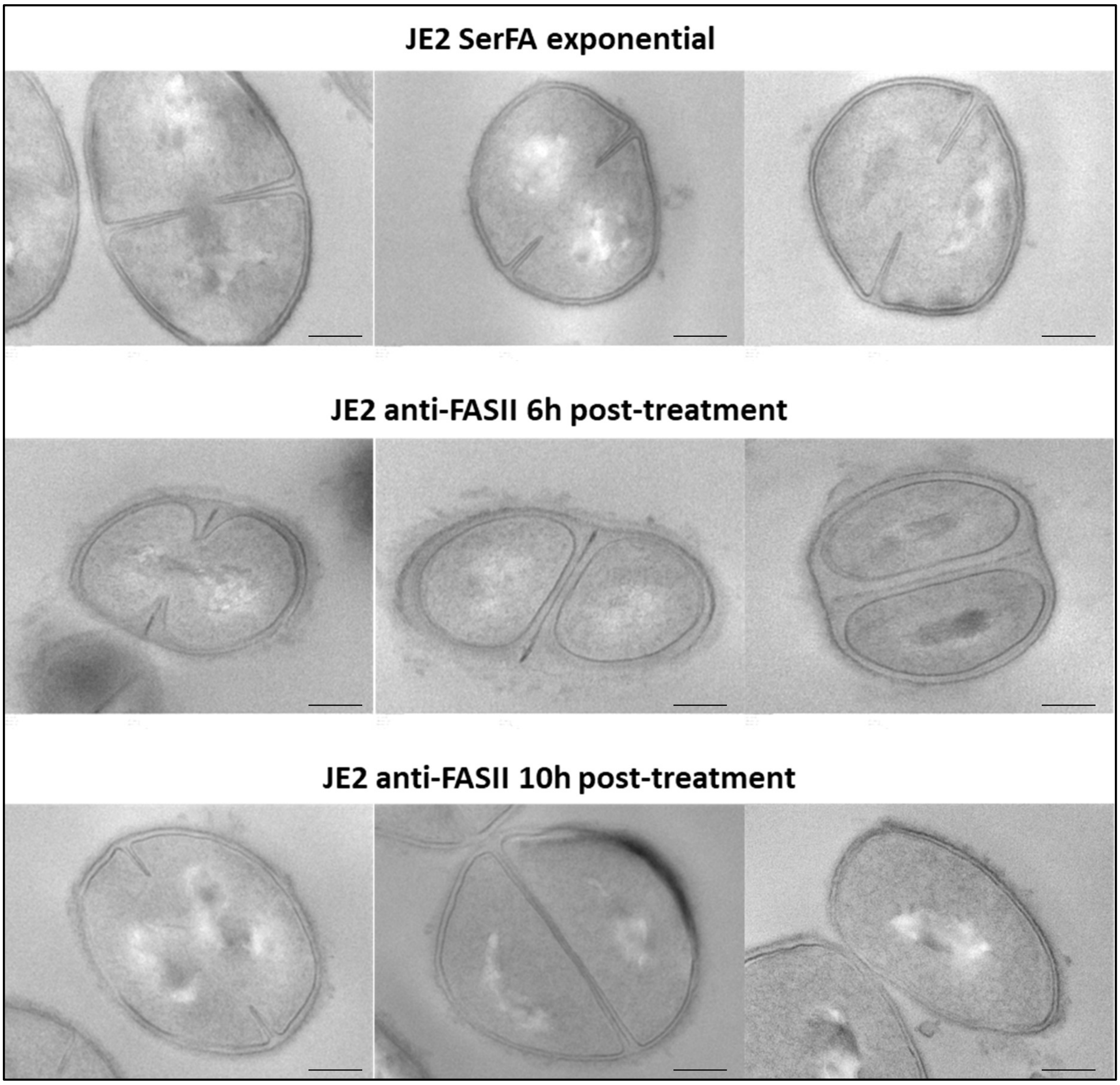


**Supplementary Fig. S2. Transmission electron microscopy of *S. aureus* JE2 in non-treated and anti-FASII-adapted growth at 6 and 10 hours.** Bar, 200 nm. Micrographs complement those shown in **Fig. 2**.

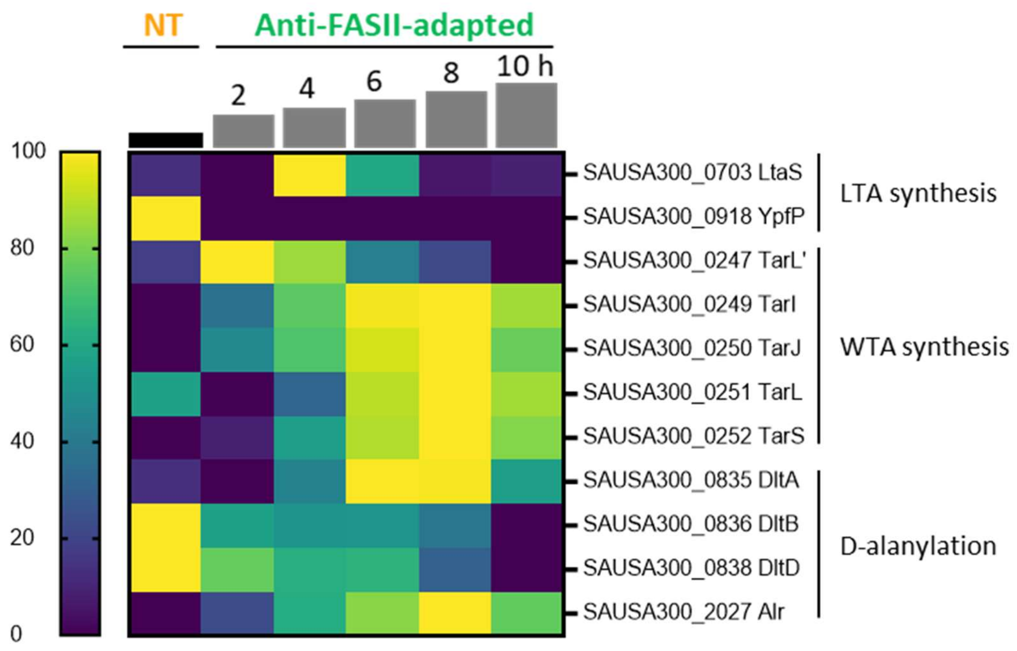


**Supplementary Fig. S3. Kinetic heatmap of *S. aureus* envelope synthesis proteins whose levels are altered during FASII$^{bypass}$.** Proteomics analyses were performed previously on *S. aureus* USA300 JE2 strain grown in SerFA, and treated or not with the FabI inhibitor triclosan (0.5 µg/ml), performed on biological quadruplicates (from [35]). Sampling times (in hours) in above steps correspond to 2, 4, 6, 8, and 10 h post anti-FASII-treatment. The heat map shows changes in detected LTA and WTA biosynthetic enzymes, and D-alanylation enzymes, which mediate decoration of both structures [75]. Changes in protein expression were determined relative to weighted values for each protein (scale at left: navy, down-represented and yellow, up-represented). WTA biosynthetic enzymes TarL' (also called TarK) and TarL are redundant [76]. Data is derived from [35].

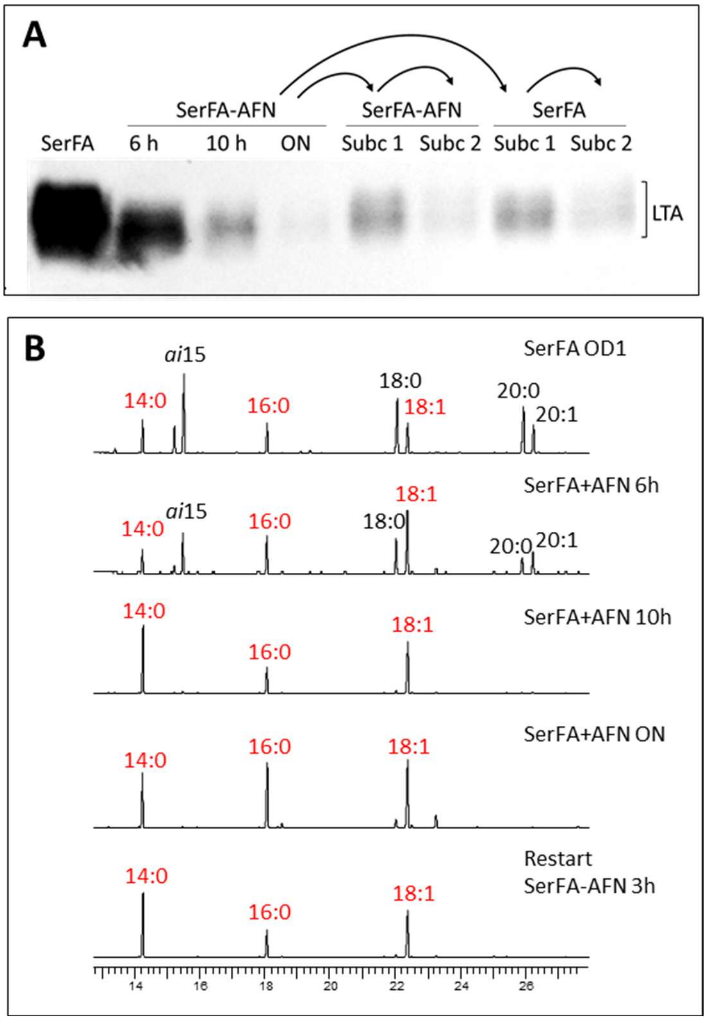


**Supplementary Fig. S4. LTA remains depleted after prolonged incubation in anti-FASII-adapted *S. aureus*. A.** *S. aureus* JE2 was grown in SerFA medium without and with AFN-1252 and samples were harvested at 6 or 10 hours, or after overnight (ON) growth. The ON anti-FASII-adapted cultures were sub-cultured (Subc 1) into fresh SerFA-AFN or SerFA medium (indicated by curved arrows) and grown 3 h. A second subculture (Subc1 to Subc 2) was prepared in the same condition. Whole cell extracts were prepared and LTA was detected by immunoblotting using anti-LTA antibody. LTA remains depleted even after long term adaptation (N=3). **B.** Samples from **A** were extracted for FA analyses at the indicated $OD_{600}$ or time. Red, exogenous FAs; black, major endogenous FAs (N=2). Supports data from **Fig. 3**.

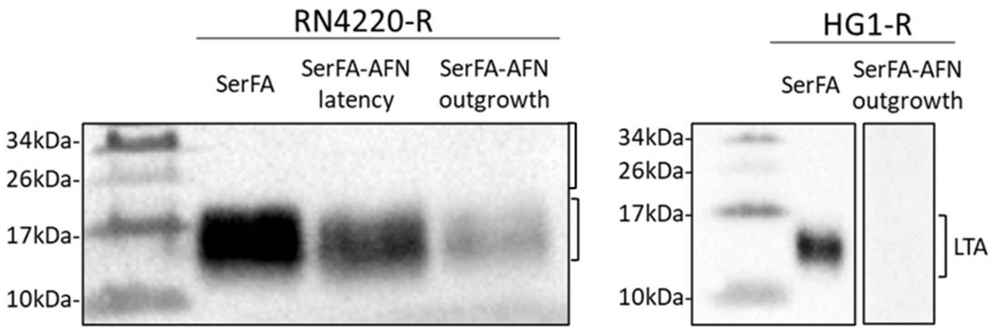


**Supplementary Fig. S5. LTA detection in *S. aureus* NCTC 8325 derivatives.** *S. aureus* RN4220 and HG001 are both from the NCTC 8325 lineage, which lacks the functional fakB1 gene required for complete exogenous FA incorporation [77]. Repair of fakB1 generated RN-R and HG1-R respectively [64]. Cultures were grown in SerFA without (non-treated, NT) and with AFN-1252 (anti-FASII-adapted, AD). LTA production was detected by immunoblotting using anti-LTA antibody (N=3). HG1-R samples are from a single gel, immunoblot, and exposure time. Supports data from **Fig. 3**.

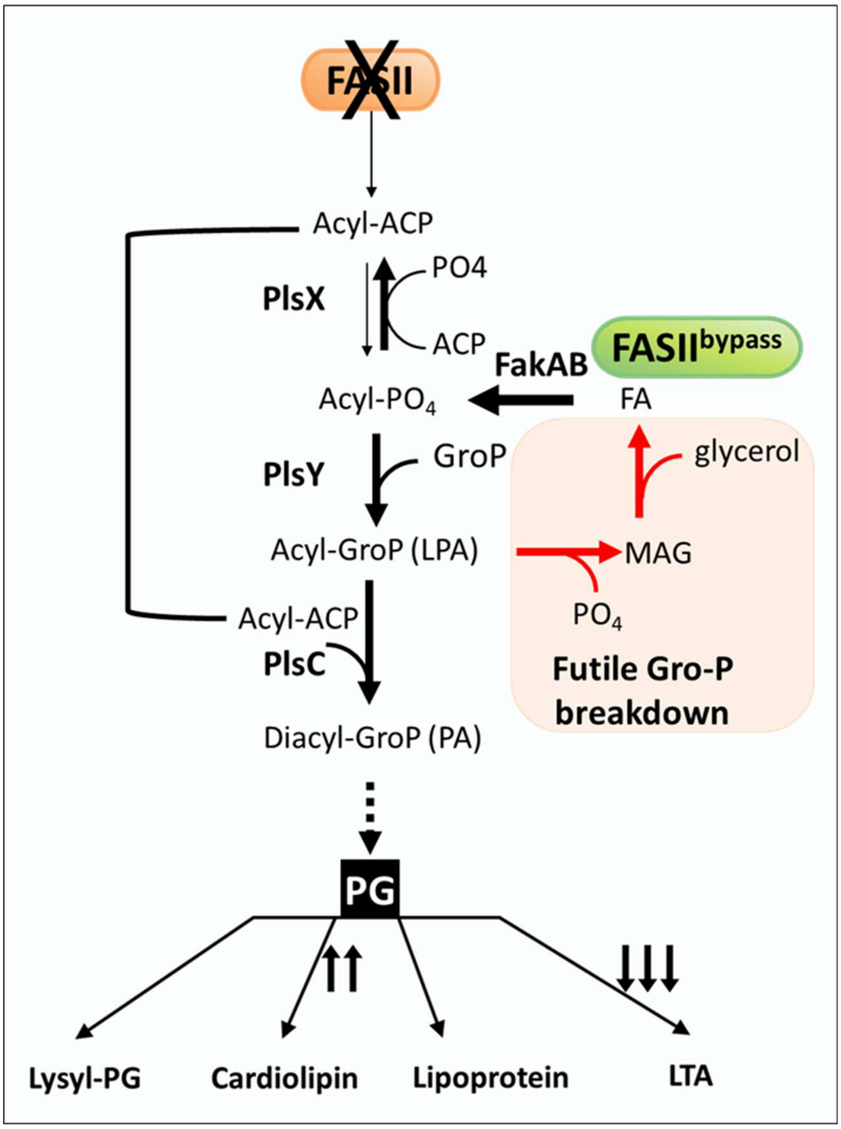


**Supplementary Fig. S6. Model linking FASII synthesis arrest to GroP metabolite depletion.** FASII inhibition due to antibiotics, mutation, and/or exogenous FA inhibition, is compensated by incorporation of exogenous FAs in membranes, schematized here for *S. aureus* [32-34, 43, 65]. Incorporation requires FakAB for FA phosphorylation [78], and reverse PlsX activity to provide acyl-ACP [33, 34]. This rerouted cycle is proposed here to desynchronize acyl-ACP and lysophosphatidic acid (LPA) availability for PlsC-mediated PA synthesis. Accumulated LPA intermediates are unstable, leading to degradation of their Gro-P moieties (salmon color zone; [62]), consistent with Gro-P depletion in anti-FASII-adapted *S. aureus* (Fig. 6). We propose that FASII blockage leads to GroP breakdown, such that the high demand for GroP from PG turnover to produce LTA cannot be met. Short arrows indicate the coordinate LTA decrease and cardiolipin increase when FASII is inhibited. Strong black arrows, favored reactions; thin black arrows, reduced or inhibited reactions; dashed arrow, multi-step reactions. Red arrows and pink zone highlight reactions leading to Gro-P depletion. The upper part of the figure applies information from [62]. The model supports results from **Fig. 6**.
