## Supplementary material for "The fatty acid synthesis pathway is a checkpoint for lipoteichoic acid synthesis in *Staphylococcus aureus*": Source file for Fig. 7A

Source data for Figure 7A.

Strains, fatty acid (FA) supplements, and addition of antimicrobial AFN-1252 (AFN) are indicated at right of each profile. BHI medium contained 10% delipidated serum. Lipidomic analyses were performed on biological duplicates (rep1, rep2).

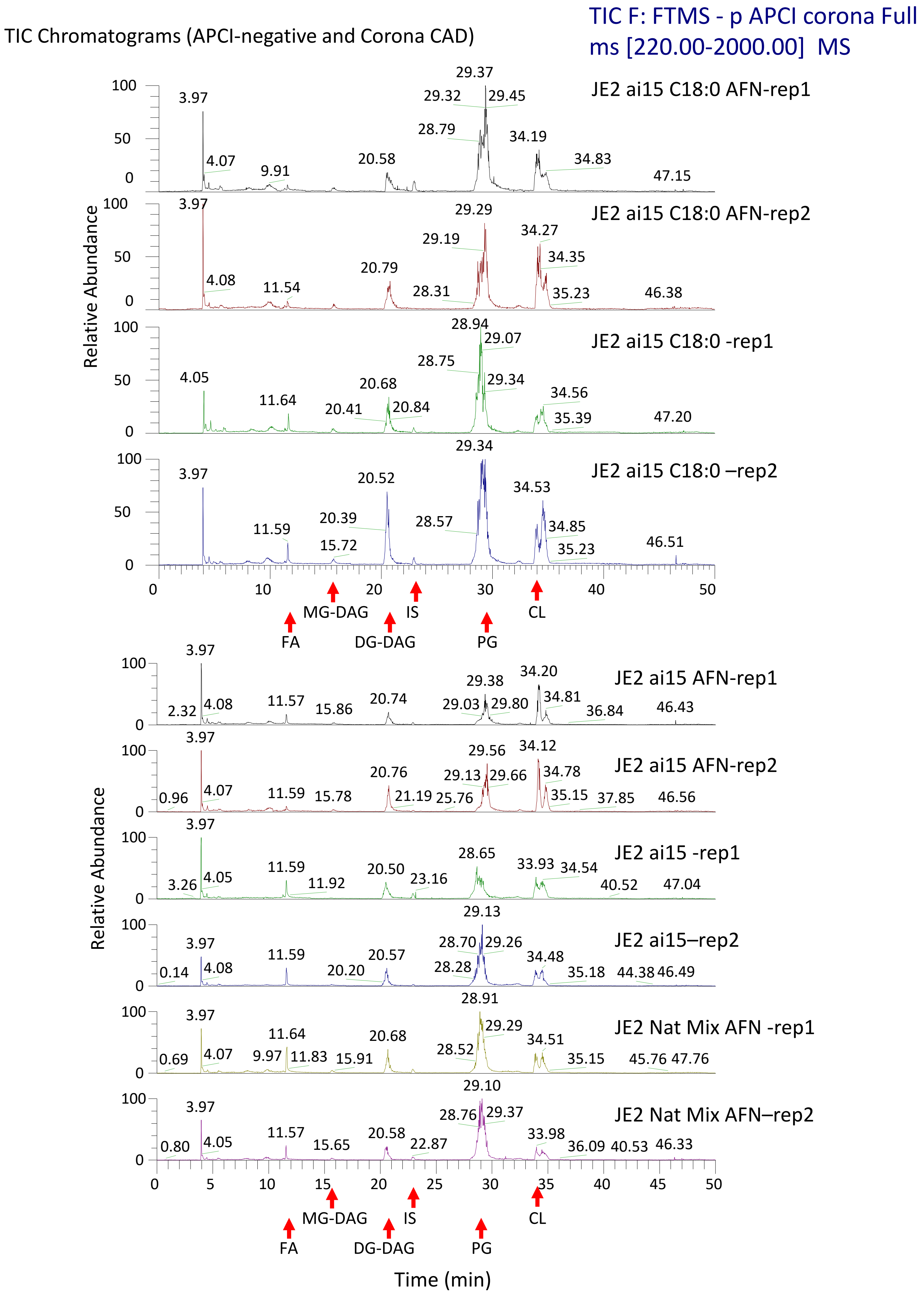

TIC Chromatograms (APCI-negative and Corona CAD)
TIC F: FTMS - p APCI corona Full ms [220.00-2000.00] MS
JE2 ai15 C18:0 AFN-rep1
JE2 ai15 C18:0 AFN-rep2
JE2 ai15 C18:0 -rep1
JE2 ai15 C18:0 –rep2
JE2 ai15 AFN-rep1
JE2 ai15 AFN-rep2
JE2 ai15 -rep1
JE2 ai15–rep2
JE2 Nat Mix AFN -rep1
JE2 Nat Mix AFN–rep2
Relative Abundance
Time (min)
FA
MG-DAG
DG-DAG
IS
PG
CL

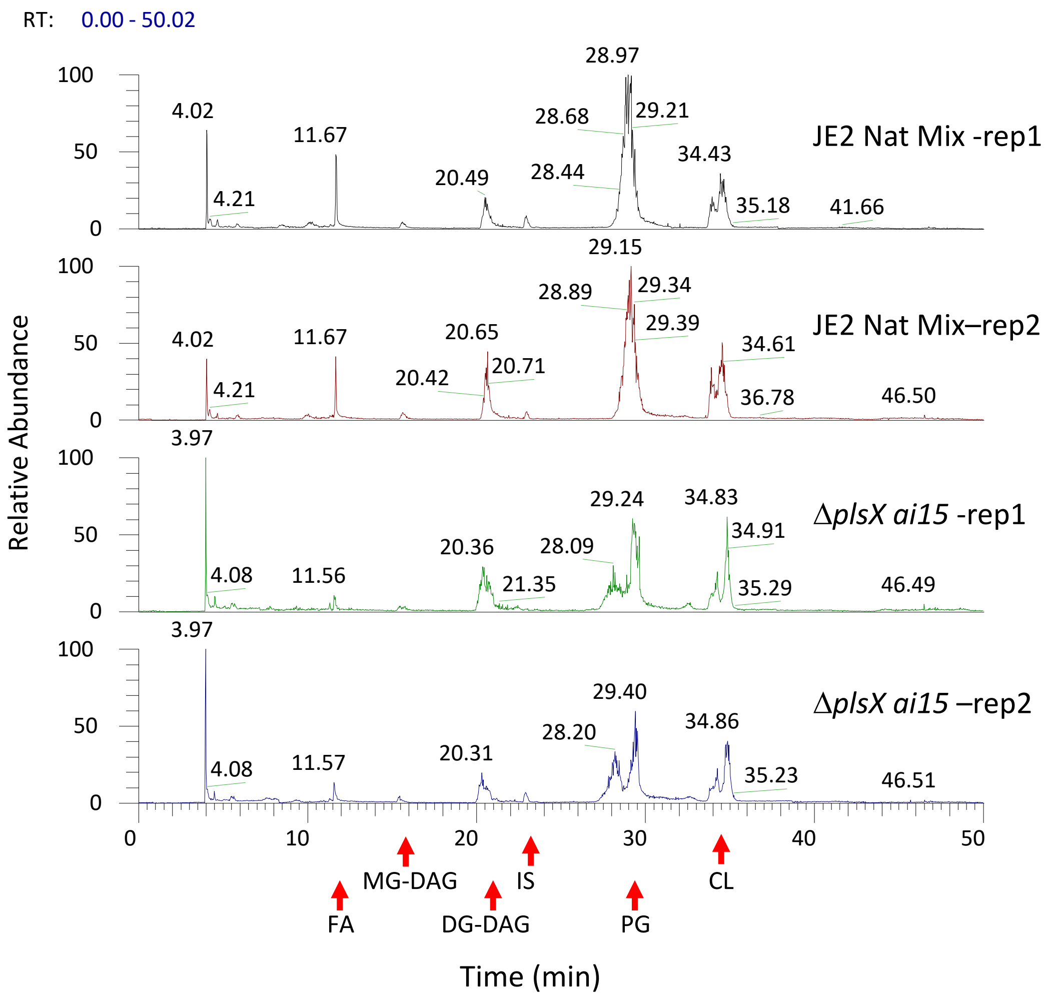


FA: free fatty acids

IS: internal standard, cyanur-phosphatidylethanolamine

MG-DAG: mono-glucosyldiacylglycerol

DG-DAG: di-glucosyldiacylglycerol

PG: phosphatidylglycerol

CL: cardiolipins

MG-DAG (15.6min)

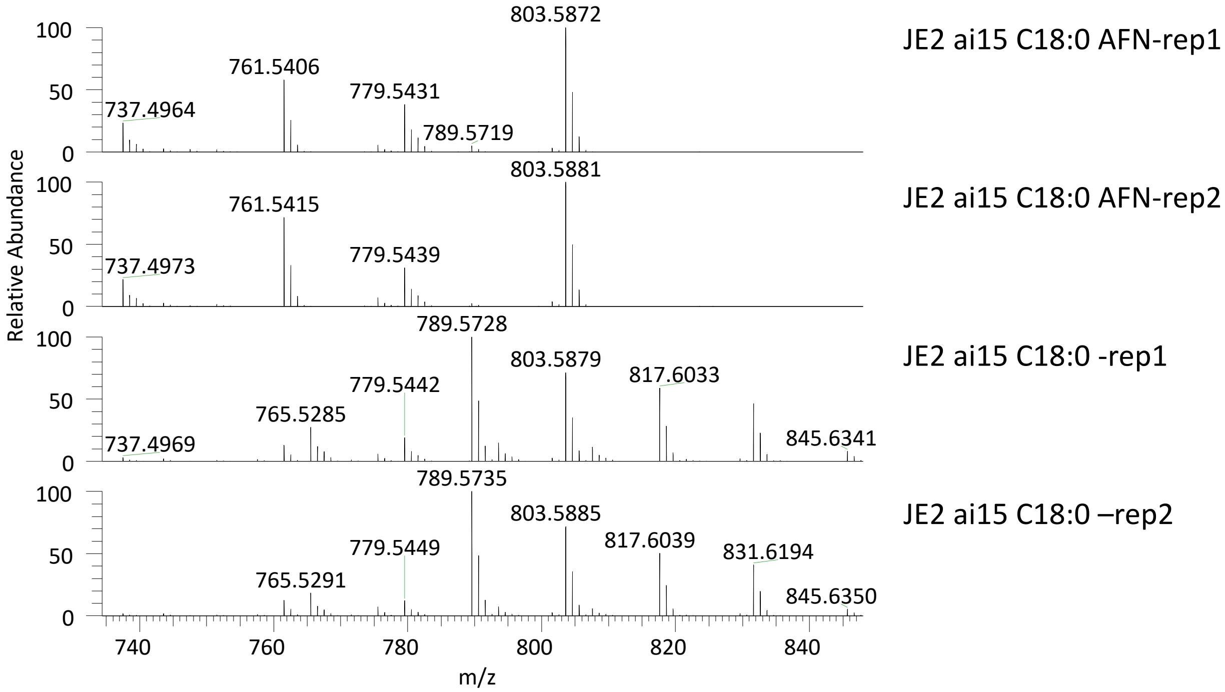


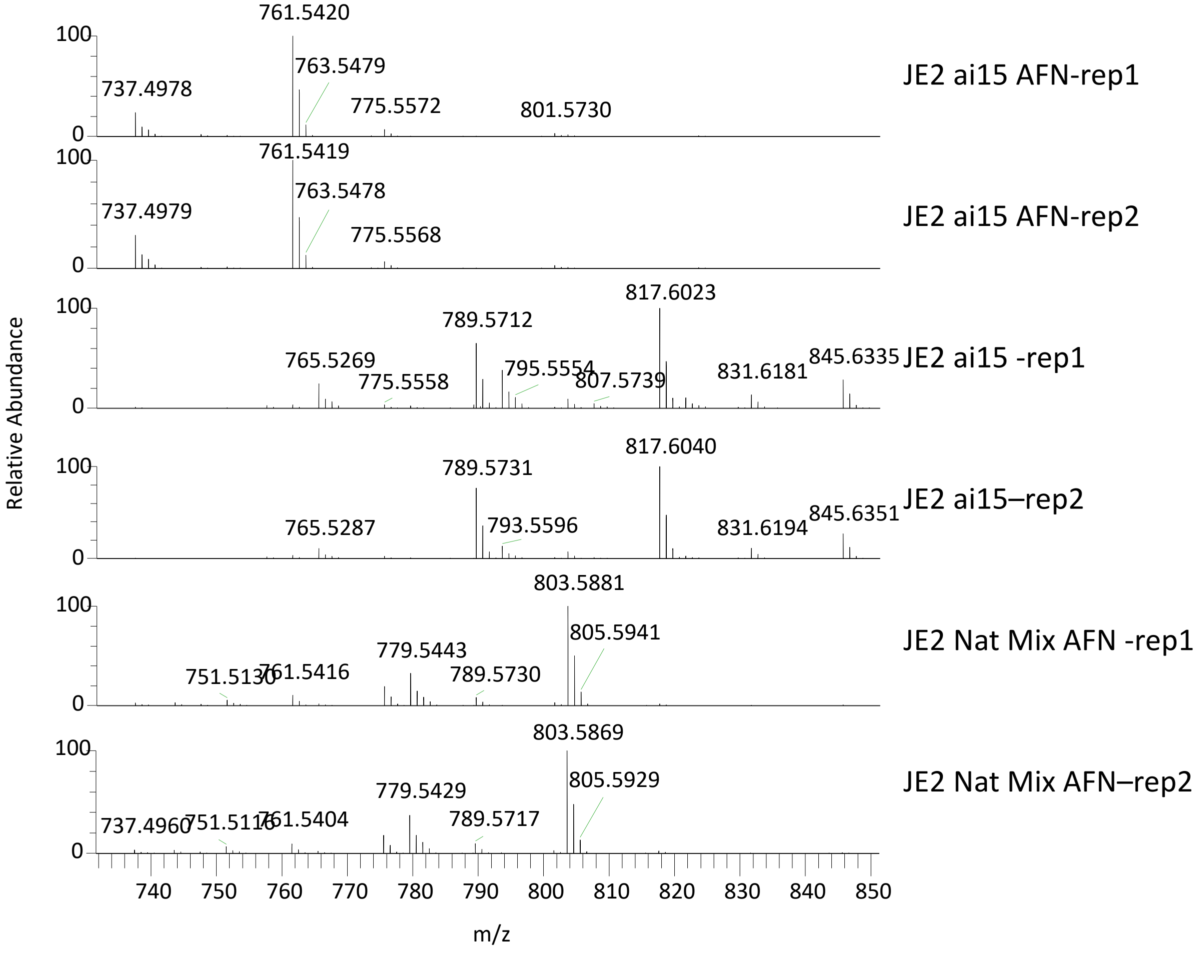

MG-DAG (15.6min)

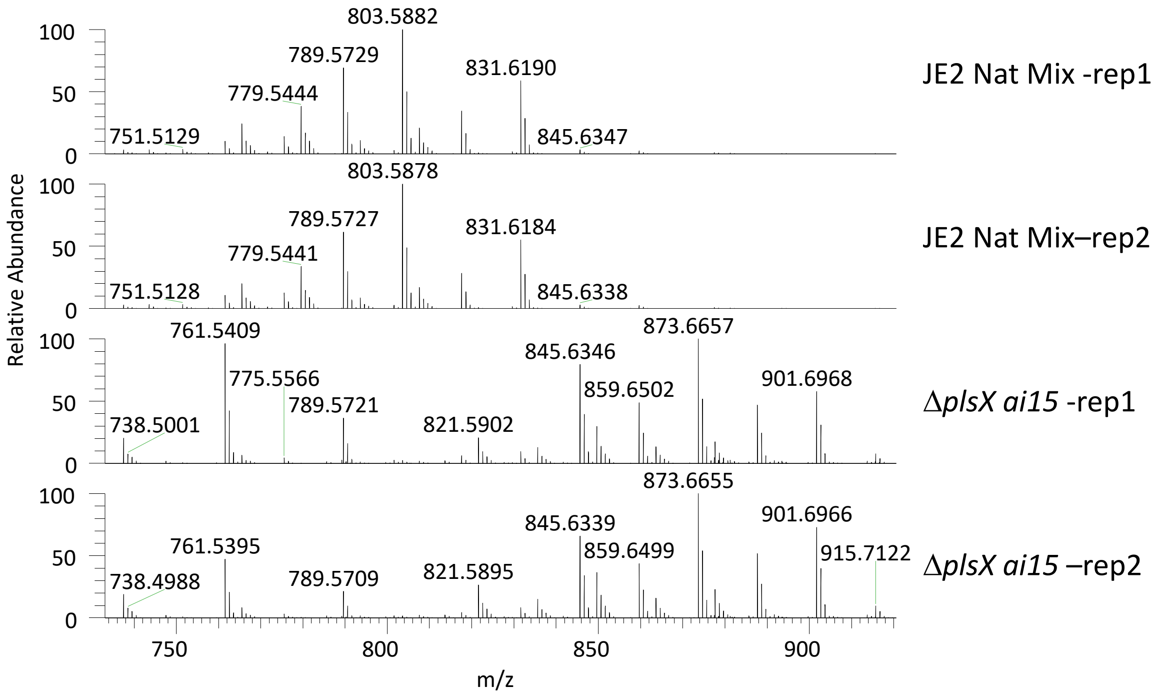

DG-DAG (20.6 min)

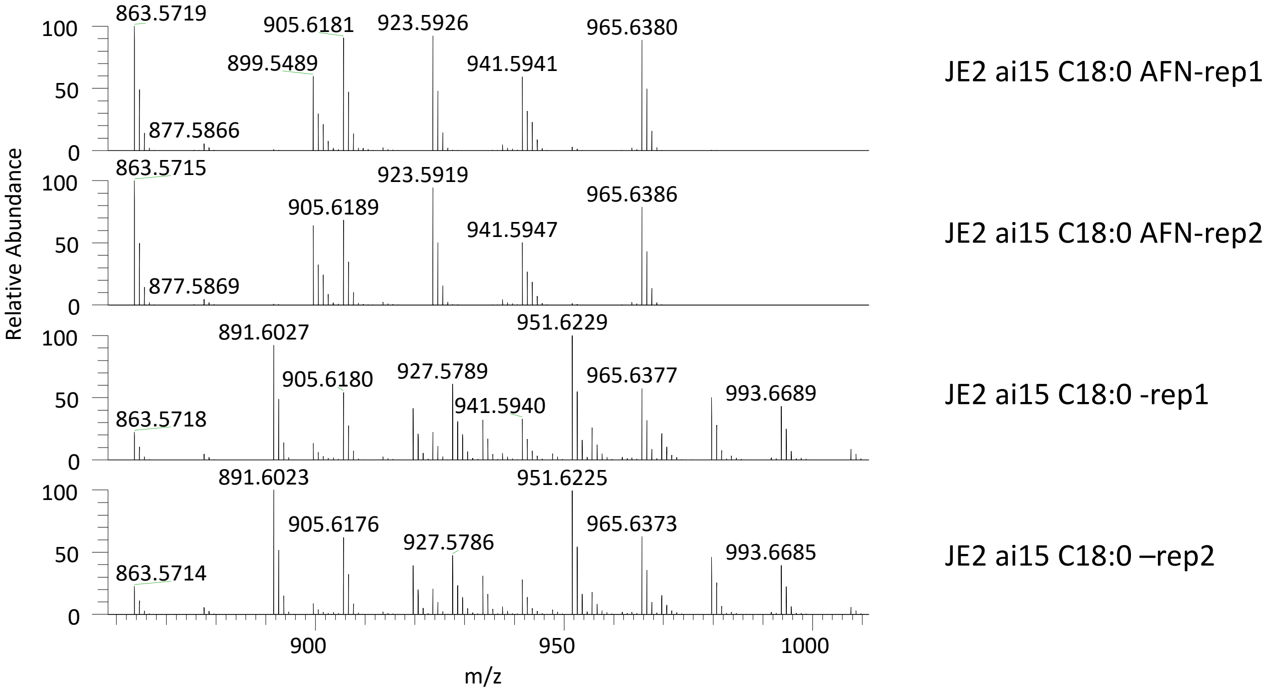


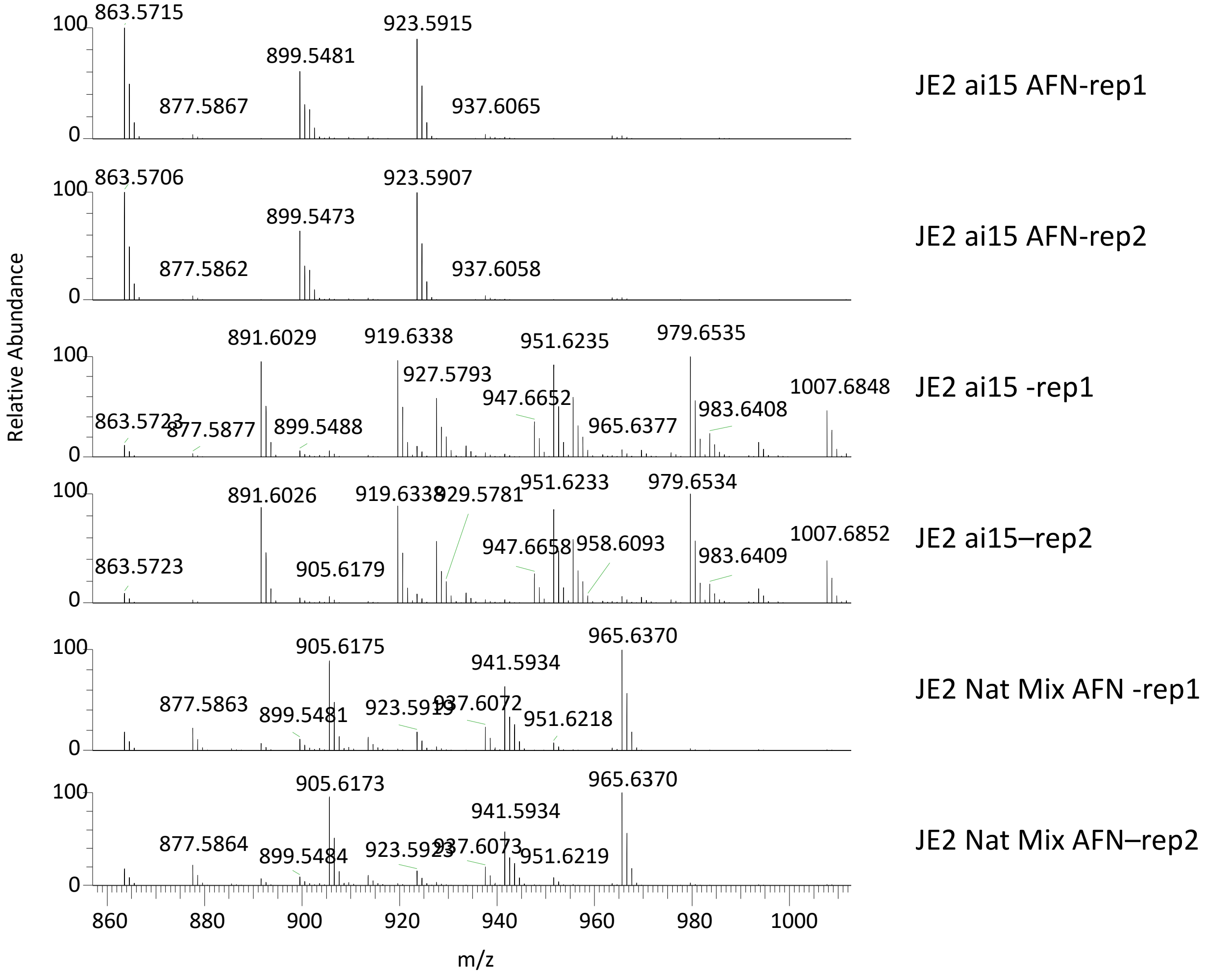

### DG-DAG (20.6 min)

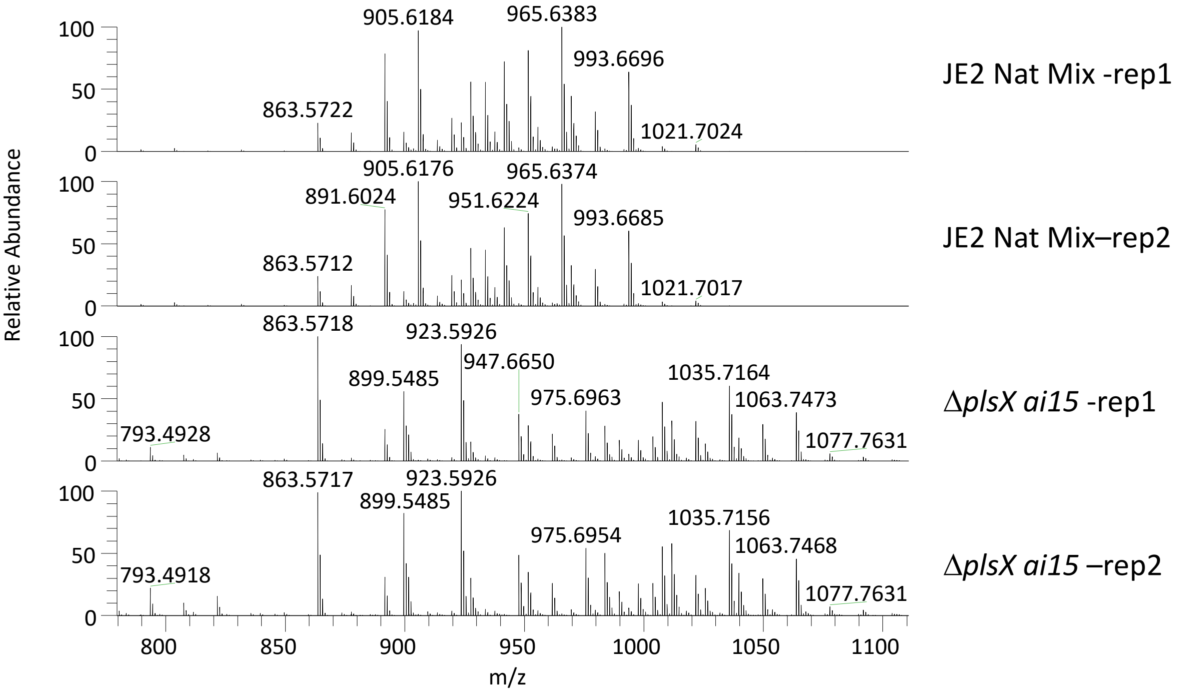

PG (28.7 min)

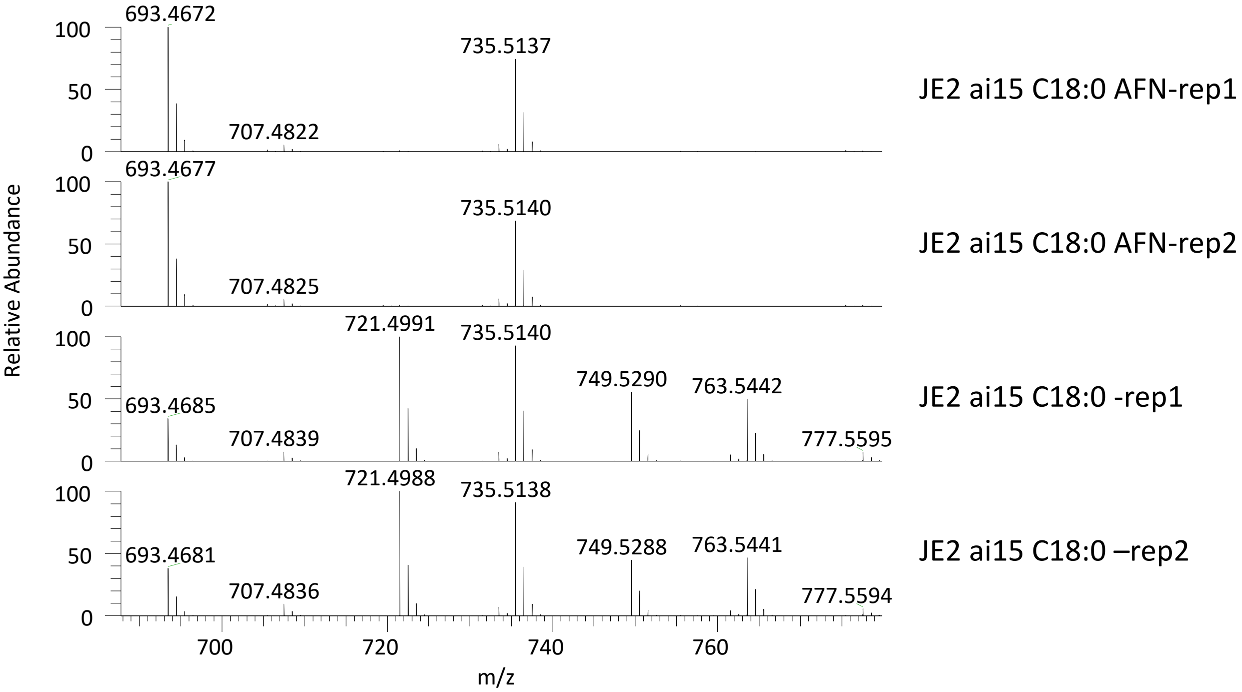
693.4672
735.5137
707.4822
JE2 ai15 C18:0 AFN-rep1
693.4677
735.5140
707.4825
JE2 ai15 C18:0 AFN-rep2
721.4991
735.5140
749.5290
763.5442
693.4685
707.4839
777.5595
JE2 ai15 C18:0 -rep1
721.4988
735.5138
749.5288
763.5441
693.4681
707.4836
777.5594
JE2 ai15 C18:0 –rep2
Relative Abundance
100
50
0
700
720
740
760
m/z

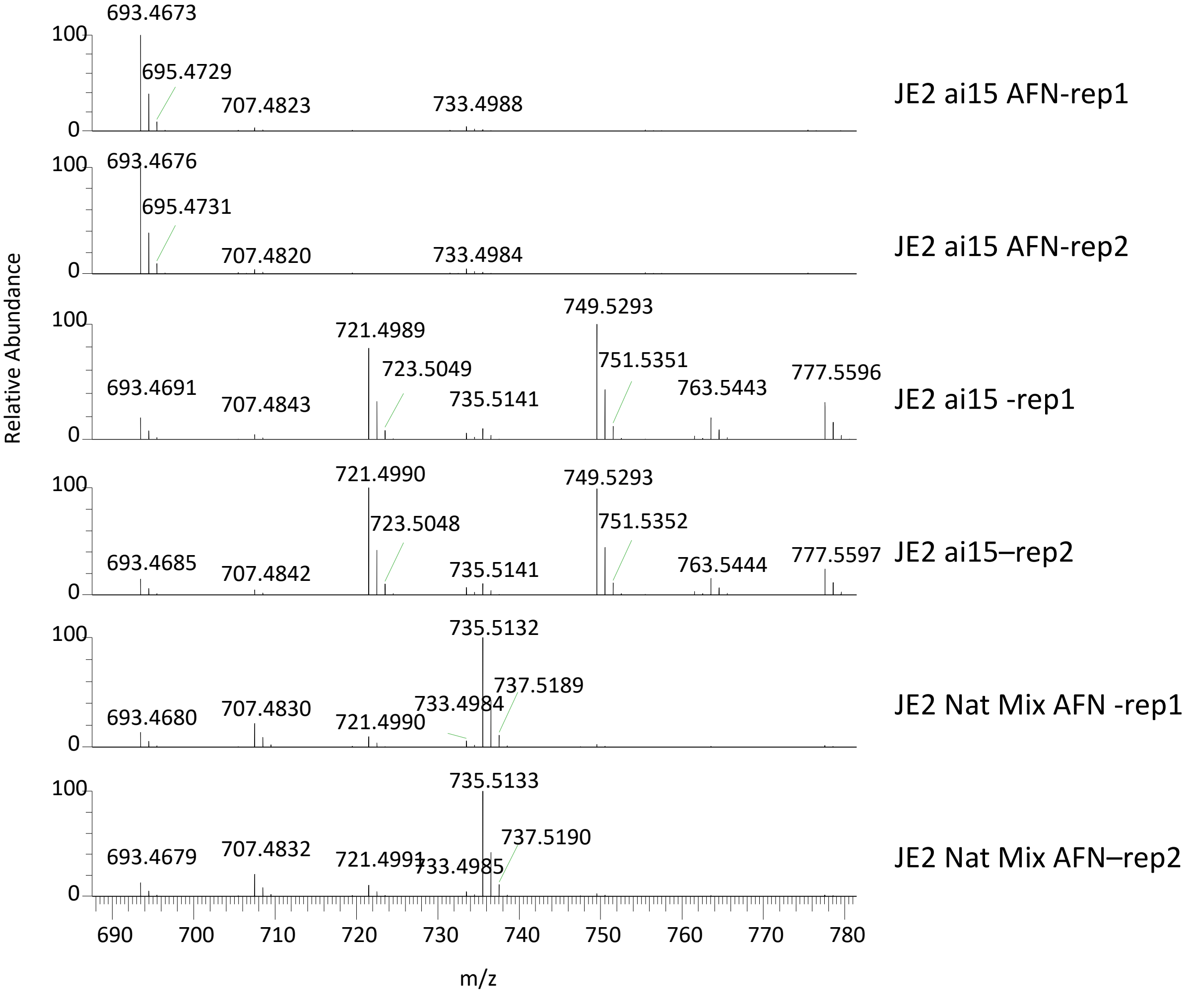
693.4673
695.4729
707.4823
733.4988
JE2 ai15 AFN-rep1
693.4676
695.4731
707.4820
733.4984
JE2 ai15 AFN-rep2
749.5293
721.4989
751.5351
723.5049
693.4691
707.4843
735.5141
763.5443
777.5596
JE2 ai15 -rep1
721.4990
749.5293
723.5048
751.5352
693.4685
707.4842
735.5141
763.5444
777.5597
JE2 ai15–rep2
735.5132
737.5189
733.4984
693.4680
707.4830
721.4990
JE2 Nat Mix AFN -rep1
735.5133
737.5190
693.4679
707.4832
721.4991
733.4985
JE2 Nat Mix AFN–rep2
Relative Abundance
100
0
690
700
710
720
730
740
750
760
770
780
m/z

### PG (28.7 min)

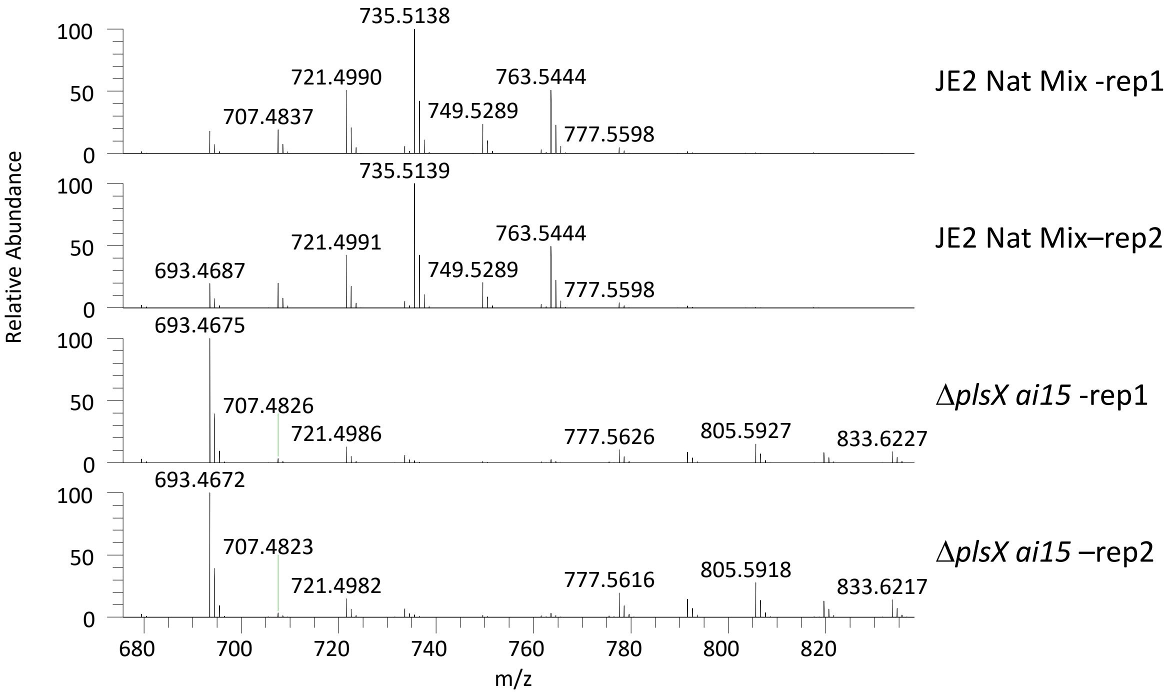
735.5138
721.4990
763.5444
707.4837
749.5289
777.5598
JE2 Nat Mix -rep1
735.5139
721.4991
763.5444
693.4687
749.5289
777.5598
JE2 Nat Mix–rep2
693.4675
707.4826
721.4986
777.5626
805.5927
833.6227
ΔplsX ai15 -rep1
693.4672
707.4823
721.4982
777.5616
805.5918
833.6217
ΔplsX ai15 –rep2
Relative Abundance
100
50
0
680
700
720
740
760
780
800
820
m/z

CL (33.8 min)

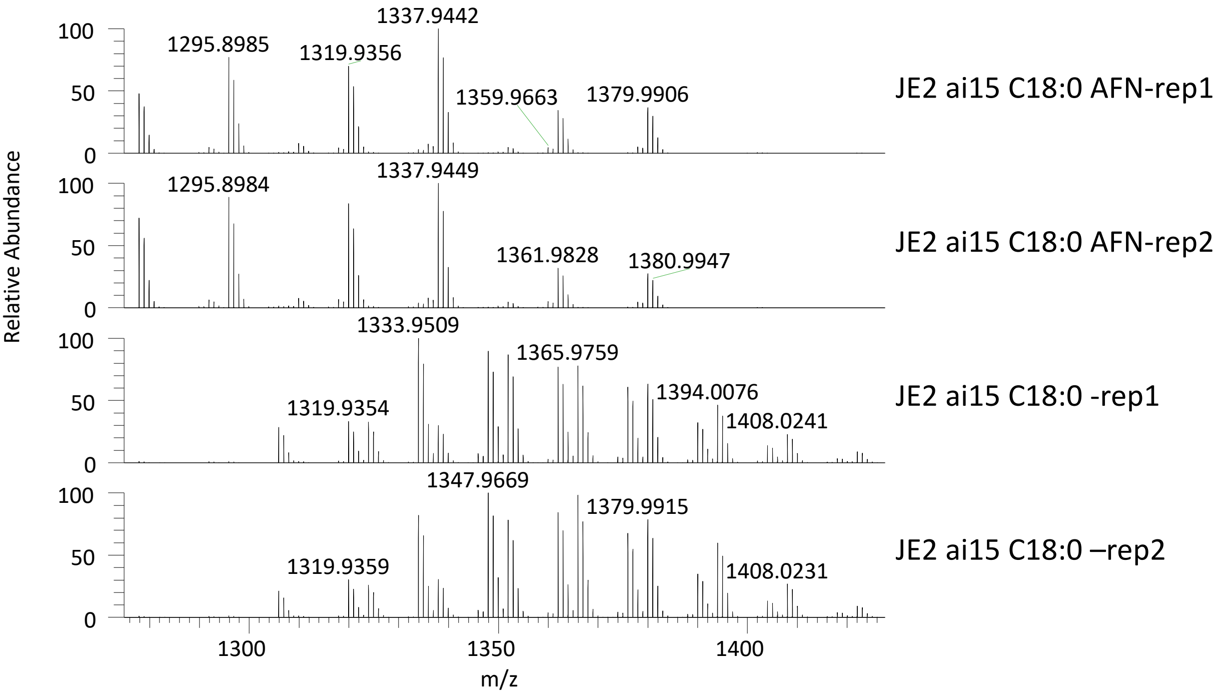
1295.8985
1319.9356
1337.9442
1359.9663
1379.9906
JE2 ai15 C18:0 AFN-rep1
1295.8984
1337.9449
1361.9828
1380.9947
JE2 ai15 C18:0 AFN-rep2
1333.9509
1365.9759
1319.9354
1394.0076
1408.0241
JE2 ai15 C18:0 -rep1
1347.9669
1379.9915
1319.9359
1408.0231
JE2 ai15 C18:0 –rep2
Relative Abundance
m/z

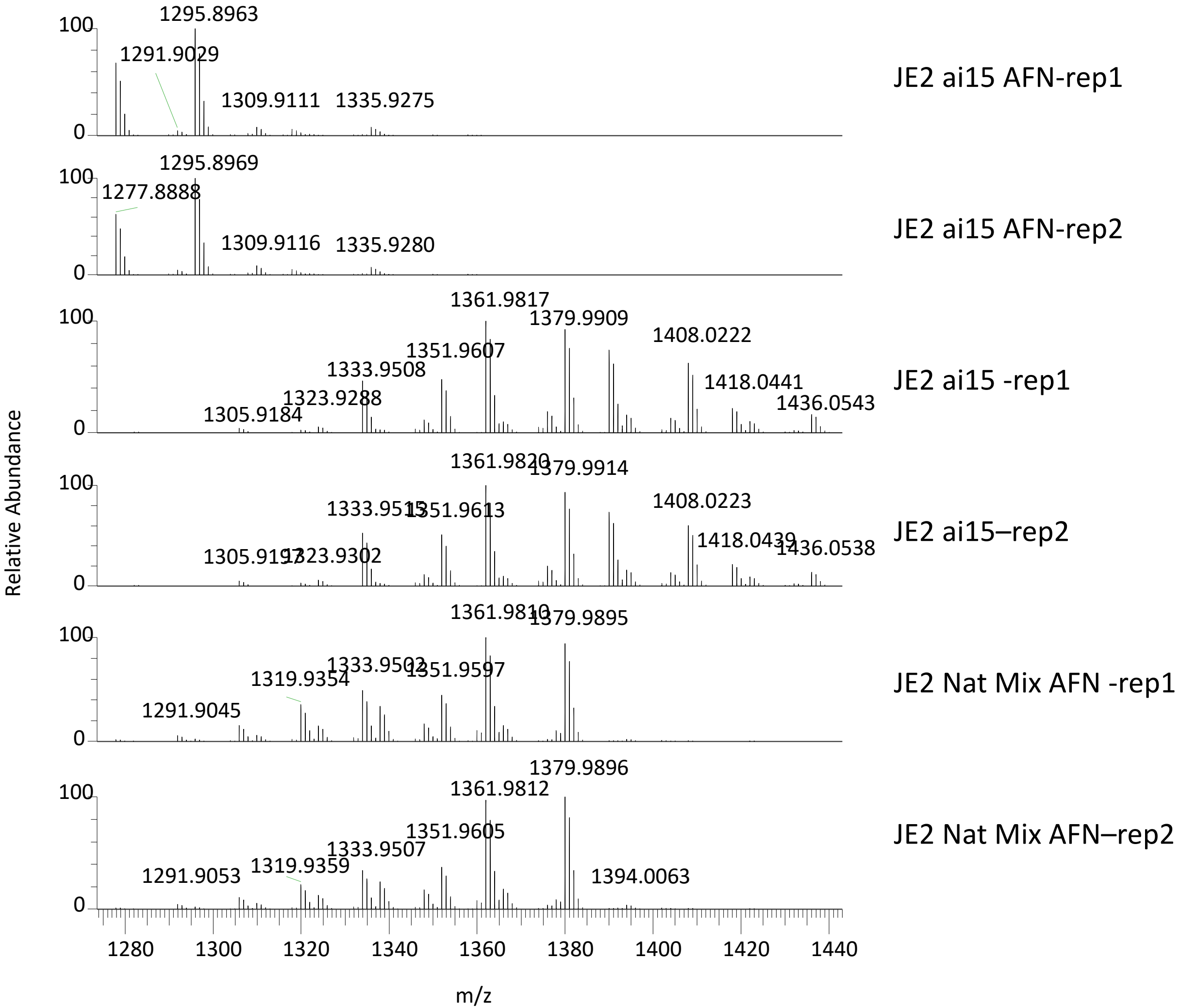
1295.8963
1291.9029
1309.9111
1335.9275
JE2 ai15 AFN-rep1
1295.8969
1277.8888
1309.9116
1335.9280
JE2 ai15 AFN-rep2
1361.9817
1379.9909
1408.0222
1351.9607
1333.9508
1323.9288
1305.9184
1418.0441
1436.0543
JE2 ai15 -rep1
1379.9914
1408.0223
1418.0439
1436.0538
JE2 ai15–rep2
1379.9895
1319.9354
1291.9045
JE2 Nat Mix AFN -rep1
1379.9896
1361.9812
1351.9605
1333.9507
1319.9359
1291.9053
1394.0063
JE2 Nat Mix AFN–rep2
Relative Abundance
m/z

CL (33.8 min)

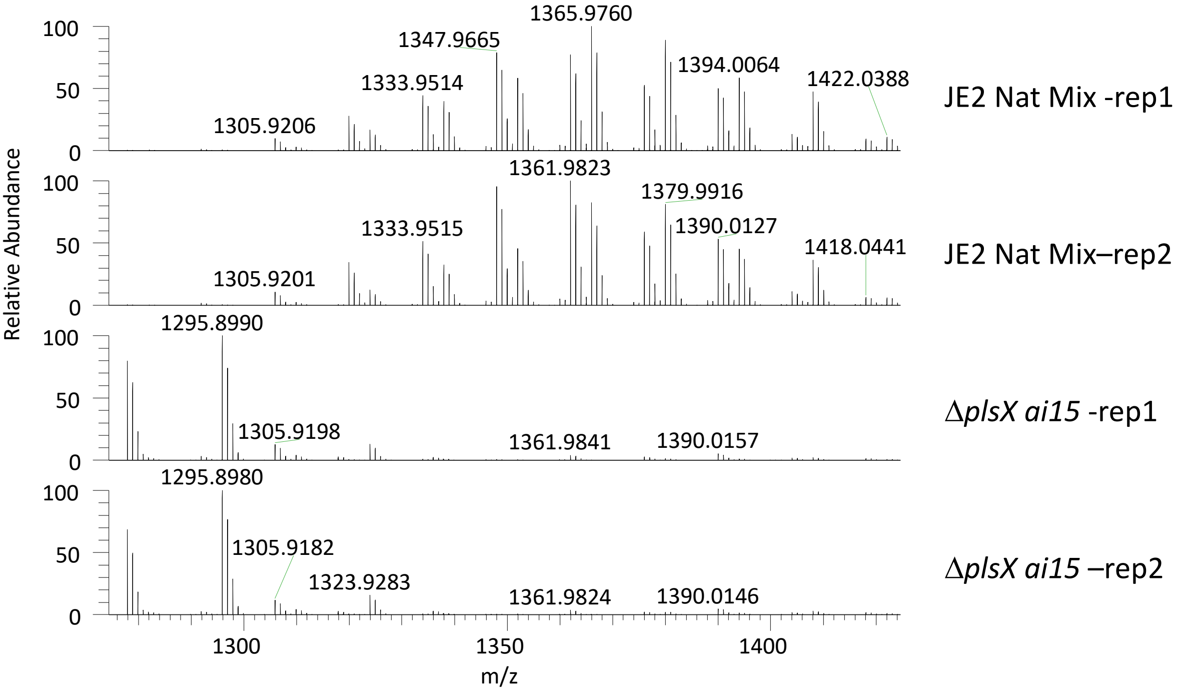

1365.9760
1347.9665
1394.0064
1333.9514
1422.0388
1305.9206
JE2 Nat Mix -rep1
1361.9823
1379.9916
1390.0127
1333.9515
1418.0441
1305.9201
JE2 Nat Mix–rep2
1295.8990
1305.9198
1361.9841
1390.0157
ΔplsX ai15 -rep1
1295.8980
1305.9182
1323.9283
1361.9824
1390.0146
ΔplsX ai15 –rep2
Relative Abundance
100
50
0
1300
1350
1400
m/z
